# Disruption of Glial Ca^2+^ Oscillations at the *Drosophila* Blood-Brain Barrier Predisposes to Seizure-Like Behavior

**DOI:** 10.1101/2020.09.16.285841

**Authors:** Shirley Weiss, Lauren C. Clamon, Julia E. Manoim, Kiel G. Ormerod, Moshe Parnas, J. Troy Littleton

## Abstract

Glia play key roles in regulating multiple aspects of neuronal development and function from invertebrates to humans. We recently found microdomain Ca^2+^ signaling in *Drosophila* cortex glia and astrocytes regulate extracellular K^+^ buffering and neurotransmitter uptake, respectively. Here we identify a role for ER store-operated Ca^2+^ entry (SOCE) in perineurial glia (PG), a distinct population that contributes to the blood-brain barrier (BBB). PG show a diverse range of Ca^2+^ oscillatory activity that varies based on their locale within the brain. Unlike cortex glia and astrocytes, PG Ca^2+^ oscillations do not require extracellular Ca^2+^ and are blocked by inhibition of SOCE or gap junctions. Disruption of these components triggers heat shock and mechanical-induced seizure-like episodes without effecting PG morphology or large molecule BBB permeability. These findings indicate SOCE-mediated Ca^2+^ oscillations in PG increase the susceptibility of seizure-like episodes in *Drosophila*, providing an additional link between glial Ca^2+^ signaling and neuronal activity.

## Introduction

Glial cells regulate multiple aspects of brain function, including synapse formation, neuronal excitability, synaptic transmission and blood flow dynamics^1^. Astrocytes, a prominent class of central nervous system (CNS) glia, modulate neuronal properties through the secretion of neuroactive agents (gliotransmission), neurotransmitter buffering and ion homeostasis, in addition to their role in synaptogenesis and blood-brain barrier function^2^. A single mammalian astrocyte forms complex interaction with various cellular counterparts via fine membranous structures collectively termed “astrocyte processes”. However, accumulating data suggest that not all processes are functionally identical^2^. The distinction between different astrocytic processes is based on morphology and supported by Ca^2+^ imaging showing astrocytes exhibit highly complex and dynamic fluctuations in intracellular Ca^2+^. These Ca^2+^ fluctuations heterogeneously distribute throughout all cellular microdomains including cell bodies, branches and endfeet that directly contact synapses and blood vessels. Although the role of glial Ca^2+^ dynamics is still being elucidated, they are hypothesized to allow astrocytes to respond to information from neighboring CNS cells and exert local modulatory control over various aspects of brain activity^2–8^.

An unresolved issue in glial biology is the contribution of distinct glial Ca^2+^ signaling components to neuronal physiology and brain function^9,10^. In addition, the signaling pathways underlying Ca^2+^ fluctuations in different cellular compartments and their implication for pathophysiology are not well understood. The controversy is often linked to the functional distinction between endoplasmic reticulum (ER)-mediated somatic Ca^2+^ oscillations and small, near-membrane microdomain Ca^2+^ oscillations. One possibility is that similar macroscopic Ca^2+^ events in astrocytes arise from diverse microscopic signaling cascades that are segregated functionally and molecularly within the cell^3,11^. An understanding of the subcellular distribution of signaling mechanisms is therefore critical to dissect how glial Ca^2+^ activity may interface with neuronal function.

Beyond glial Ca^2+^ signaling, glia also contribute to the blood-brain barrier (BBB). The BBB forms the functional metabolic separation between the CNS and circulating blood, controlling permeability of nutrients and other molecules in and out of the CNS that is necessary for brain homeostasis. The mammalian BBB consists of endothelial cells that line blood vessels, connected by tight junctions that prevent paracellular diffusion^12^. Interactions of the endothelial sheet with other cell types that reside in close proximity to the vessels such as astrocytes, pericytes, neurons, and microglia shape the homeostatic function of the BBB^12^. In *Drosophila*, the CNS is separated from the surrounding hemolymph, the insect “blood”, by a barrier that is structurally and functionally similar to the mammalian BBB (Figure 1A,^13–15^). The *Drosophila* BBB covers the entire CNS with a flattened cell sheet consisting of cellular layers formed by two classes of glia, an outer layer of perineurial glia (PG) and an inner layer of subperineurial glia (SPG)^16^. A dense extracellular matrix called the neural lamella is secreted by the PG and contributes to the BBB. The barrier function and prevention paracellular diffusion of ions and small molecules in *Drosophila* is largely mediated through septate junctions formed by adjacent SPG cells (Figure 1A,^15,17–20^). Transport of molecules across the *Drosophila* BBB occurs via membrane transport protein mechanisms in SPG, similar to the mammalian endothelial BBB^13,21^. The outermost PG layer of the BBB, together with the neural lamella, contribute to barrier selectivity against larger molecules such as proteins^15^, and is thought to serve as support for the SPG and provide structural roles^16^.

**Figure 1:**
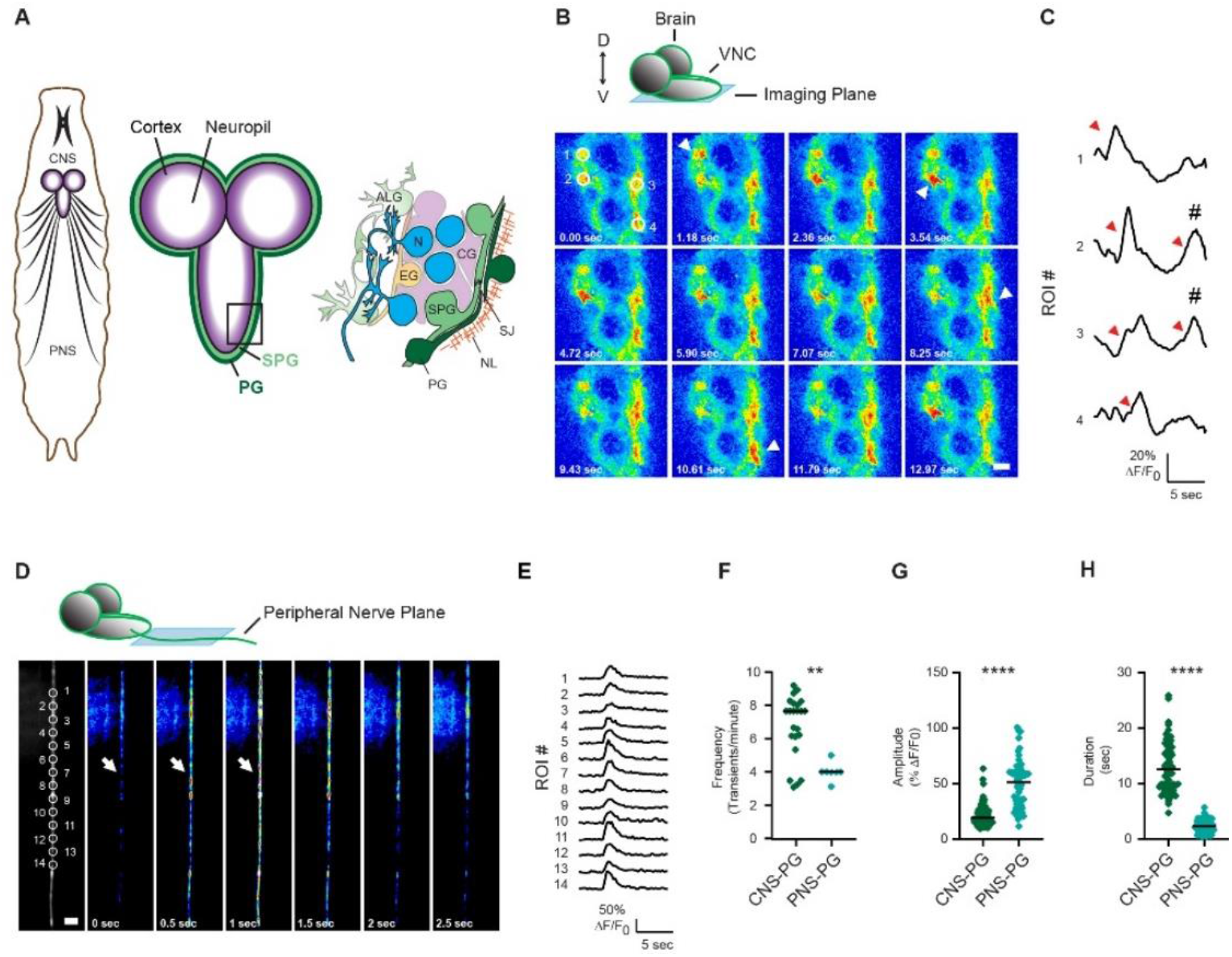
Dynamic Ca^2+^ Transients Occur in Perineural Glia In Vivo. (A) Schematic representation of the larval nervous system, showing five glial subtypes that occupy the CNS: perineurial glia (PG), subperineurial glia (SPG), cortex glia (CG), ensheathing glia (EG) and astrocyte-like glia (ALG). Neurons (N), septate junctions (SJ) and the neural lamina (NL) are also shown. (**B-H**) Ca^2+^ imaging of PG>myr::GCaMP6s in live, non-dissected 2^nd^ instar wildtype *Drosophila* larvae. (*B*) Time-lapse image series of perineural glial Ca^2+^ at the ventral surface of the VNC (CNS-PG). Arrowheads mark the peaks of Ca^2+^ transients shown in (C). Scale bar, 20 μcm. Schematic representation of the *Drosophila* larval brain (top) shows the relative field of view at the ventral surface of the VNC (light blue). The dorsal-ventral (D-V) axes is shown. (**C**) Mean myr::GCaMP6s fluorescence (% Δ*F/F*0) reveals Ca^2+^ rise spreads through adjacent ROIs (marked with red arrowheads). Respective ROIs are shown in panel A. (*D*) Time-lapse image series of perineural glial Ca^2+^ signals in an abdominal segment peripheral nerve (PNC-PG). Arrows mark local Ca^2+^ elevations that are synchronized between adjacent ROIs, as seen in panel D. Greyscale panel shows the average projection of all images in the series. Scale bar, 20 μm. (*E*) Mean myr::GCaMP6s fluorescence (% Δ*F/F*0) reveals synchronous Ca^2+^ elevation in adjacent ROIs from images in panel D. (*F-H*) Comparisons of Ca^2+^ transient characteristics in PG cells of the VNC (CNS-PG) or enwrapping peripheral nerves (PNS-PG). (*F*) Transient frequencies are significantly lower in PNS-PG (transients/minute, p<0.01, n=23 ROIs/4 VNCs, n=5 nerves/5 animals). (*G*) Transient amplitudes are significantly higher in PNS-PG (% Δ*F/F*0, p<0.0001, n=99 transients/23 ROIs/4 VNCs, n=65 transients/5 nerves/5 animals). (*H*) Transient durations are significantly shorter for PNS-PG (seconds, p<0.0001, n=61 transients/23 ROIs/4 VNCs, n=61 transients/5 nerves/5 animals). **=P < 0.01, ****=P < 0.0001, Student’s t-test.

Several *Drosophila* glial subtypes have been found to influence neuronal function via distinct mechanisms downstream of their glial Ca^2+^ dynamics. Astrocytic Ca^2+^ regulates neurotransmitter uptake^22^ and secretion of neuromodulators^23^, while disruption of Ca^2+^ signaling in *Drosophila* cortex glia impairs K^+^ buffering capacity^24^. Synchronized Ca^2+^ waves in *Drosophila* SPG cells control nutrient-dependent reactivation of neural stem cells and subsequent brain growth^25,26^. As multiple components of the vertebrate BBB show dynamic fluctuations in [Ca^2+^]_i_, we sought to explore whether *Drosophila* PG cells also exhibit fluctuations in [Ca^2+^]_i_, and if this source of Ca^2+^ signaling regulates neural activity. By performing detailed Ca^2+^ imaging, we find that PG cells exhibit complex and dynamic Ca^2+^ activity that differs between PG cells that occupy distinct brain domains. In addition, we find that these Ca^2+^ dynamics are independent of extracellular Ca^2+^, originate from internal ER stores and spread as waves through gap junctions. In a genetic screen for glial pathways that modulate neuronal excitability ^24^, we found that pan-glial knockdown of dStim, the *Drosophila* homologue of mammalian Stromal Interaction Molecule 1, leads to severe heat-shock (HS) and mechanically-induced seizure-like episodes (hereafter referred to as seizures, see methods for definition). Here we demonstrate that dStim is primarily required in PG cells for seizure induction. dStim regulates the store-operated Ca^2+^ entry (SOCE) pathway, in which the plasma membrane Ca^2+^ channel, Orai, is gated by the ER-localized Ca^2+^ sensor, Stim, leading to Ca^2+^ influx and restoration of ER Ca^2+^ stores. Similar to Stim knockdowns, disruption of Orai function in PG cells also disrupts their Ca^2+^ dynamics and increases seizure susceptibility. Inhibiting propagation of Ca^2+^ activity within the PG sheet through gap junctions recapitulates the behavioral phenotype of SOCE knockdown. While Orai and Stim are expressed in mammalian glial cells, the role of SOCE in glial function is poorly understood.

## Results

### Dynamic Ca^2+^ Transients Occur in Perineurial Glia *In Vivo*

In vertebrates, astrocytic Ca^2+^ fluctuations heterogeneously distribute between different cellular microdomains. These Ca^2+^ fluctuations occur in processes that contact synapses and neuronal cell bodies, as well as in endfeet that directly contact blood vessels. Astrocyte-like glial subtypes in *Drosophila* exhibit intracellular elevations in Ca^2+^, similar to mammalian astrocytes. As multiple components in the vertebrate BBB, including astrocytes, show fluctuations in Ca^2+^, we explored whether perineurial glia (PG) that contribute to the *Drosophila* BBB exhibit fluctuations in Ca^2+^ and how they might affect brain function.

To explore *in vivo* Ca^2+^ dynamics in PG cells, we expressed a myristoylated variant of Ca^2+^-sensitive GFP (myr::GCaMP6s) to monitor Ca^2+^ in fine processes^24,27^ with the PG driver GMR85G01-gal4^28^ (hereafter referred to as PG-gal4). We first performed imaging experiments in live, undissected 2^nd^ instar larvae as previously described^27^. PG expression of myr::GCaMP6s revealed highly complex and dynamic Ca^2+^ transients in PG cells that enwrap the ventral nerve cord (VNC) and peripheral nerves (Figure 1). The observed Ca^2+^ transients have different characteristics depending on the spatial domain that PG cells enwrap in the nervous system. PG cells that enwrap the VNC (CNS-PG) show slow local elevations in Ca^2+^ along the VNC (Figure 1B, C). These events recur frequently in the same regions, occasionally spreading as waves between neighboring cells (Figure 1C, arrowheads, F), suggesting adjacent PG cells may laterally transfer information through Ca^2+^ waves. The duration of CNS-PG Ca^2+^ transients was 12.96±0.59 seconds and exhibited a mean *Δ*F/F_*0*_ of 21.08±0.9% (Figures 1G, H). In contrast, PG cells that enwrap peripheral nerves (PNS-PG) show fast and synchronous elevations in Ca^2+^ along the whole imaging field (Figure 1D, E). These transients recur frequently (~4 events/min, Figure 1F), with a duration of 2.32±0.142 seconds and a mean *Δ*F/F_*0*_ of 49.11±2.61% (Figure 1G, H). These data indicate PG cells display dynamic Ca^2+^ signaling *in vivo*, with PGs in the CNS or PNS displaying distinct patterns of Ca^2+^ oscillatory activity.

### Characterization of PG Ca^2+^ Activity Reveals Unique Signatures in Cells that Occupy Different CNS Territories

To further explore PG Ca^2+^ signaling, we developed a semi-dissected preparation to image the larval CNS. GCaMP6s expressing dorsal PG cells in the CNS were imaged in HL3.1 external saline^29,30^. To minimize muscle contractions, we first performed imaging recordings in external solution containing no added Ca^2+^ (i.e. nominal [Ca^2+^]_out_). Surprisingly, PG cells in the VNC (VNC-PG) exhibited robust population-wide waves under these conditions (Video 1A, Figures 2A, S1A-C), suggesting PG Ca^2+^ signaling relies on intracellular Ca^2+^ stores rather than extracellular influx. PG cells at the VNC exhibited robust, slow Ca^2+^ transients with a mean *Δ*F/F_*0*_ of 53.63±1.48% (Figure S1D) and a duration of 20.20±0.95 seconds (Figure S1E). Ca^2+^ waves at the VNC can be synchronized across neighboring regions in the PG sheet (Figure S1B, C). This synchronous Ca^2+^ activity likely represents the activity of multiple PG cells. To characterize the activity of single cells, we co-expressed myr::GCaMP6s and mCherry.nls in PG cells (Video 1B,C, Figure 2A, B) and assigned regions of interest (ROI) to single cells based on nuclear labeling. Single cells at the VNC showed Ca^2+^ activity patterns in which cells alternate between rhythmic activity and silent periods (Figure 2B). This activity pattern can be synchronized between adjacent cells (Figure 2B, lower superimposition trace). Ca^2+^ transients in single VNC-PG cells during rhythmic activity periods exhibited large amplitudes, with a mean *Δ*F/F_*0*_ of 91.10±3.28% (Figure 2C, D and H). The mean duration of single VNC-PG transients was 20.42±0.56 seconds (Figure 2I), similar to what was measured with randomly assigned ROIs (Figure S1E).

**Figure 2:**
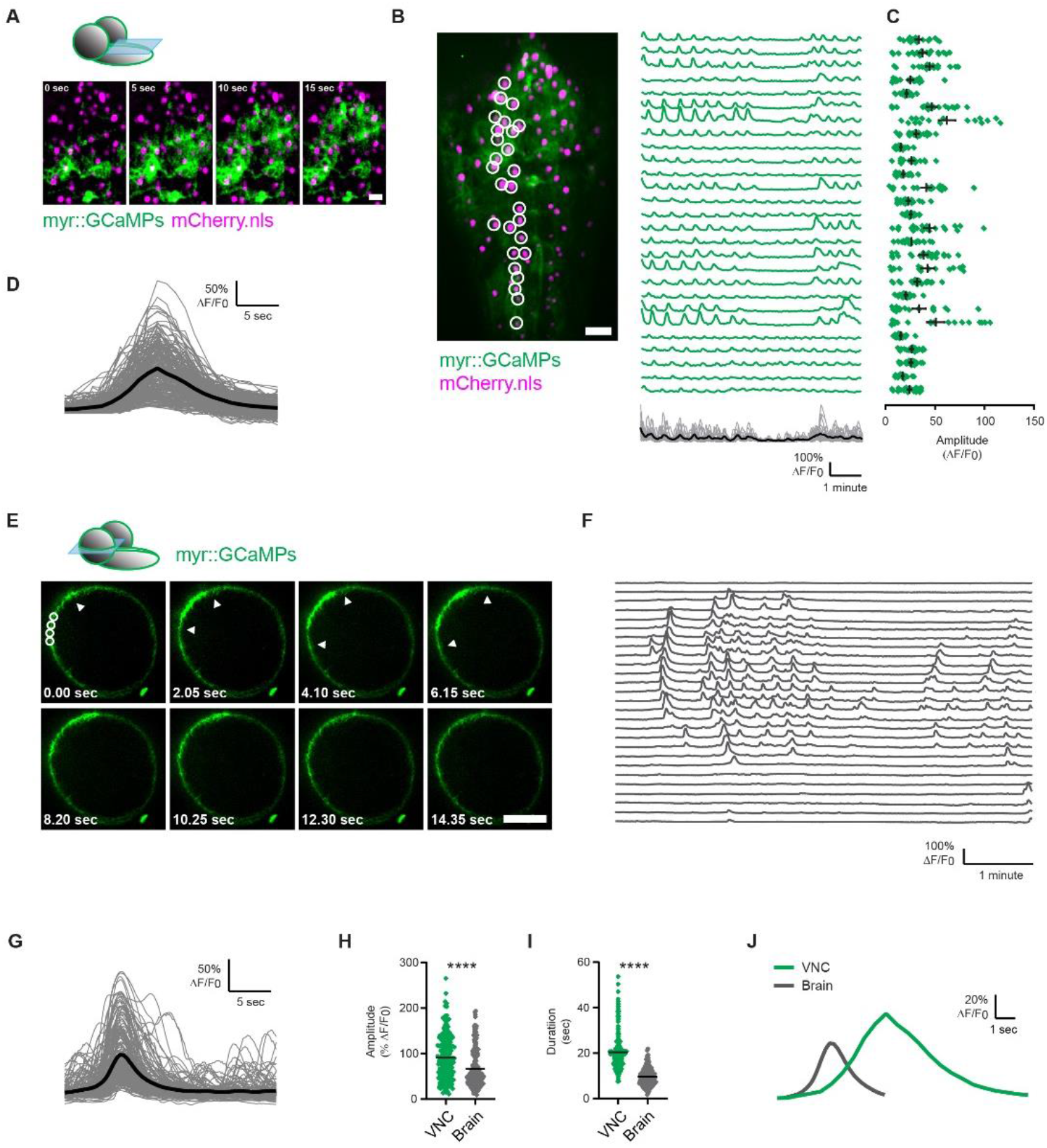
Characterization of PG Ca^2+^ Reveals Distinct Signatures in Cells that Occupy Different CNS Territories. Imaging of PG>myr::GCaMP6s in dissected 3_rd_ instar wildtype *Drosophila* larvae. (**A-D**) Ca^2+^ activity of single PG cells co-expressing myr::GCaMP6s and nuclear-localized mCherry (magenta: mCherry.nls, PG nuclei; green: myr:GCAMP6s, PG membrane). (**A**) Representative time-lapse image series of PG Ca^2+^ at the dorsal surface of the VNC (VNC-PG), showing a Ca^2+^ wave spreading through multiple adjacent PG cells (See Video 1). Scale bar, 20 μm. (**B**) Analysis of single cell Ca^2+^ activity. ROIs were assigned to single cells by mCherry.nls as shown (left). Representative traces of mean myr::GCaMP6s fluorescence (% Δ*F/F*_0_) of single cells reveal dynamic Ca^2+^ activity that is synchronized between a large number of cells across the VNC. Bottom trace shows superimposition of all traces (n=27, grey) with the mean highlighted in black. Scale bar, 20 μm. (**C**) Ca^2+^ transient amplitudes for the traces in (B) are shown. (**D**) Superimposition of VNC-PG Ca^2+^ transients (gray) and their mean (black; n=222 individual transients/60 ROIs/3 animals). Only 4 representative ROIs are shown. (**E**) Time-lapse image series of Ca^2+^ imaging in PG that enwrap a brain hemisphere (brain-PG). Mid-section is shown (See Video 2). Arrowheads mark the lateral spread of the Ca^2+^ wave. (F) Representative traces of mean myr::GCaMP6s fluorescence (% Δ*F/F*_0_) show dynamic Ca^2+^ activity that spreads across multiple adjacent cells (see Figure S2J for demonstration of Ca^2+^ wave spread). (**G**) Superimposition of single brain-PG Ca^2+^ transients (gray) and their mean (black; n=205 transients/60 ROIs/3 animals). (**H-I**) Comparisons of Ca^2+^ transient characteristics in VNC-PG and brain-PG (N>200 transients/60 ROIs/3 animals/CNS region). VNC-PG transients show significantly higher amplitudes (p<0.0001) (**G**) and longer durations (p<0.0001) (*E*). (**I**) Superimposition of the means of VNC-PG and brain-PG Ca^2+^ transients (from Figures 2D and 2G). ****=p<0.0001, Student’s t-test.

In contrast to the slow waves observed in VNC-PG, PG cells on the surface of the brain (Brain-PG) showed fast, rhythmic, asynchronous activity (Video 2A and Figures S1F-H). Transients that occurred between Ca^2+^ waves (see below) recurred frequently in the same region (~4 events/min, Figure S1G) with a mean *ΔF/F_0_* of 29.56±0.7% (Figure S1H). Imaging Ca^2+^ from a mid-section through a brain hemisphere revealed that brain-PG exhibit robust Ca^2+^ waves that spread through large areas of the PG sheet (Figures 2E-G, S1I and Video 2B, note that sporadic, single cell events are also visible). Relative to VNC-PG, Ca^2+^ transients observed in brain-PG exhibited smaller amplitudes, with a mean *Δ*F/F_*0*_ of 66.38±2.91% (Figure 2H) and a shorter duration with a mean duration of 9.71±0.26 seconds (Figures 2I, J). PNS-PG cells did not show Ca^2+^ oscillations under these experimental conditions, suggesting that PNS-PG signaling relies more on extracellular Ca^2+^. Together, these data indicate that PG cells show complex and diverse Ca^2+^ activity patterns based on their location, suggesting functional diversity within the PG cell population.

### Ca^2+^ Transients in Perineurial Glia Originate from Internal ER Ca^2+^ Stores

The occurrence of PG Ca^2+^ transients in a low-Ca^2+^ external solution suggests that PG Ca^2+^ signaling is likely to rely on internal ER Ca^2+^ stores. To test this hypothesis, we repeated the Ca^2+^ imaging experiments in an external solution containing 10 μM Thapsigargin (Tg, see Methods) to pharmacologically inhibit restoration of ER Ca^2+^ stores. Under these conditions PG Ca^2+^ signaling in both the VNC and the brain is almost completely abolished (Figures 3A and Video 2C). These data indicate that pharmacological inhibition of ER Ca^2+^ stores with Tg significantly reduces PG Ca^2+^ signaling, supporting the model that PG Ca^2+^ activity relays solely on Ca^2+^ signals that originate from internal ER Ca^2+^ stores.

**Figure 3:**
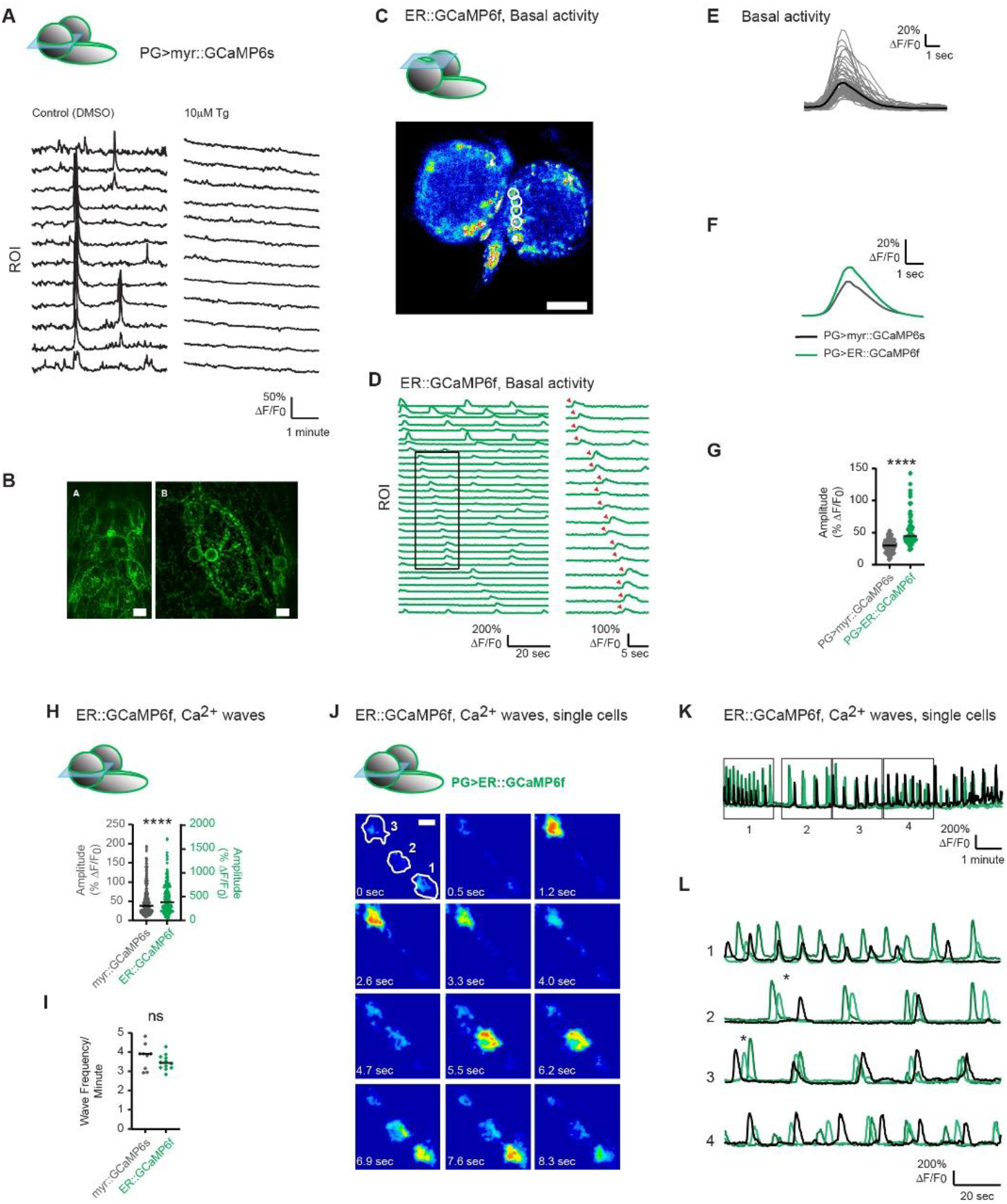
ER Originated Ca^2+^ Waves in PG Cells Display Striking Complexity. (**A**) Pharmacologically inhibiting ER Ca^2+^ signaling in dissected 3^rd^ instar wildtype *Drosophila* larvae. Representative traces of mean myr:: GCaMP6s fluorescence (% Δ*F/F*_0_) of randomly assigned ROIs in brain-PG show dynamic Ca^2+^ activity under control conditions (1% DMSO) that is completely abolished when samples are incubated in 10μM Thapsigargin (Tg) for 2 minutes prior to imaging. Rapid decrease in basal myr:GCaMP6s fluorescence prevented automated detection of small events. (**B**) Live imaging of the cellular localization of ER::GCaMP6f at the dorsal surface of the VNC in 3^rd^ instar *Drosophila* larva. Z-stack projection of PG on the dorsal surface of the VNC reveal that ER accumulates around the nuclei and cell periphery. (Scale bars; left: 20 μm, right: 5 μm). (**C-L**) Imaging of PG>ER::GCaMP6s in dissected 3^rd^ instar wildtype *Drosophila* larvae. (**C-G**) Single cells at the dorsal surface of a brain hemisphere. Scale bar, 50 μm. (**C**) Heat map summarizing spontaneous PG activity on the surface of the brain determined by GCaMP6s imaging. ROIs were assigned to single cells based on activity maps. (**D**) Representative traces of mean ER::GCaMP6s fluorescence (% Δ*F/F*_0_) of single cells show dynamic Ca^2+^ activity that is asynchronized between neighboring cells. Inset on the right shows lateral spread of Ca^2+^ elevations through adjacent cells. (**E**) Superimposition of single brain-PG ER-originated Ca^2+^ transients (gray) and their mean (black; n=73 transients/50 ROIs/3 animals, only events between waves were analyzed). (**F**) Superimposition of the means of myr:GcaMP6s and ER::GCaMP6f Ca^2+^ transients in brain-PG. (**G**) Comparison of isolated Ca^2+^ transient amplitudes (% Δ*F/F*_0_) recorded with myr:GcaMP6s and ER::GCaMP6f shows significant increase (p<0.0001) in ER::GCaMP6f (n=78 transients, median is presented) recorded events relative to myr:GcaMP6s (n=110 transients, median is presented). (**H**) Comparison of Ca^2+^ wave amplitudes (% Δ*F/F*_0_) recorded with myr:GcaMP6s and ER::GCaMP6f shows significant increase (p<0.0001) in ER::GCaMP6f recorded wave amplitude relative to myr:GcaMP6s (n=156 transients for myr:GCaMP6s and n=73 transients for ER::GCaMP6f, medians are presented).(**I**) Comparison of the frequency of Ca^2+^ waves (event/minute) recorded with myr:GcaMP6s (n=9) and ER::GCaMP6f (n=12) shows no significant difference (medians are presented). (**J**) Time-lapse image series of PG Ca^2+^ with ER::GCaMP6f. Enlarged view of 3 neighboring brain-PG cells is shown (see Video 3). Scale bar, 5 μm. (**K**) Representative traces of mean ER::GCaMP6s fluorescence (% Δ*F/F*_0_) of single cells show dynamic and complex Ca^2+^ activity with dynamic frequencies that are synchronized between cells. (**L**) Two-minute insets of the traces from (J) show complex cell-to-cell Ca^2+^ spread.

### Characterization of ER-Originated Ca^2+^ Signals in Perineurial Cells Reveals Complex Intra- and Inter-Cellular Signaling

Astrocytes exhibit highly complex and dynamic fluctuations in Ca^2+^ that vary between different cellular compartments. To explore ER-originated Ca^2+^ signals in PG cells, we generated a transgenic *Drosophila* line expressing GCaMP6f tethered to the external surface of the ER^31^ (ER::GCaMP6f, see Methods). Examination of the cellular localization of ER::GCaMP6f in PG in dorsal VNC revealed flattened, tile-like morphology (Figure 3B), with ER that fills the entire cellular volume and accumulates around the nucleus and in periphery of the cell where PG cell-to-cell contacts are formed (Figure 3B, right). This expansive network is consistent with a role for ER signaling in cell-to-cell communication. Imaging basal Ca^2+^ activity (activity that occurs between Ca^2+^ waves) of single PG cells that tile the brain hemispheres (brain-PG) revealed fast, rhythmic, asynchronous activity similar to that observed with myr:GCaMP6s (compare Figures 3C-E and S1F-H). These Ca^2+^ transients recurred frequently in the same regions (~3 events/min, Figure 3D) and showed a significant increase in amplitude (relative to events recorded with myr::GCaMP6s) with a mean *Δ*F/F_*0*_ of 53.02±2.9% (Figures 3F-G). Even in the absence of large amplitude waves, some ER-originated Ca^2+^ elevations appear to pass through multiple adjacent cells (Figure 3D, inset, red arrowheads). The similar wave forms of the transients measured with ER::GCaMP6f and myr::GCaMP6s (Figure 3F), together with similar frequencies, indicate that the majority of events in PG cells originate from ER Ca^2+^ stores.

Similar to what was observed with myr:GCaMP6s, hemisphere mid-section Ca^2+^ imaging with ER::GCaMP6f revealed robust Ca^2+^ waves that spread through multiple neighboring PG cells (Video 3). Interestingly, the amplitudes recorded during waves are significantly larger than those of isolated transients that occur during basal, inter-wave signaling, with a mean amplitude of 461%±20.9% (Figure 3G, H). This observation suggests an enhancement mechanism that increases the Ca^2+^ signal while it spreads through the PG sheet. In addition, as expected from the localization of the sensor at the Ca^2+^ source, the amplitudes recorded during waves using ER::GCaMP6f are significantly larger than those observed with the membrane-bound myr::GCaMP6s sensor (Figure 3H). The wave frequencies recorded with the two sensors are similar (~4 transients/ minute, Figure 3I), providing additional support that all events observed in PG cells originate from internal ER Ca^2+^ stores. Finally, close examination of the relations between the activity of adjacent cells reveals striking complexity (Figure 3J-L, Video 3): synchronized changes in transient frequencies (Figure 3K and insets 1-4 in Figure 3L), Ca^2+^ waves spreading without a preferred direction (* in Figure 3L) and highly variable latencies of the Ca^2+^ spread between neighboring cells. Together, these data indicate PG cells exhibit complex and diverse ER originated Ca^2+^ activity that propagates through adjacent cells to form robust Ca^2+^ waves that spread over long distances in the *Drosophila* nervous system.

### Genetic Screening Reveals Glial Knockdown of dStim Increases Seizure Susceptibility in *Drosophila*

We recently found that chronic Ca^2+^ increase in *Drosophila* cortex glia predispose animals to stimulation-induced seizures^24^, while acute increases in intracellular Ca^2+^ in *Drosophila* astrocyte-like glia drives neuronal silencing^22^. To identify glial signaling pathways that may modulate neuronal excitability, we performed a genetic screen using the pan-glial driver repo-gal4 to drive expression of RNA interference (RNAi) targeting ~850 genes encoding membrane receptors, secreted ligands, ion channels and transporters, vesicular trafficking proteins and known cellular Ca^2+^ homeostasis and Ca^2+^ signaling pathway components^24^. This screen revealed that pan-glial knockdown of dStim led to severe HS-induced seizures-like episodes (Video 4, middle and Figure 4A, S2A), similar to those we previously identified in *NCKX^zyd^* (*zyd*) mutants that disrupt a Ca^2+^ ion exchanger in cortex glia_24,27_. Pan-glial knockdown of dStim (repo>dStim^RNAi^) on the *zyd* mutation background lead to ~30% of flies showing room-temperature seizures and enhanced HS-induced seizures (Figure S2A), indicating the two manipulations are likely disrupting independent pathways to enhance seizure susceptibility. Recordings of the motor central pattern generator (CPG) output at the larval neuromuscular junction (NMJ) demonstrated that 3^rd^ instar repo>dStim^RNAi^ larvae lose normal rhythmic firing at 38°C and instead display continuous neuronal firing, as observed in *zyd* mutants (Figure S2B). The seizure phenotype that resulted from dStim knockdown was similar when dStim was targeted using four additional partially overlapping dStim RNAi constructs (Figure S2C, see Methods). All five dStim^RNAi^ lines also showed seizures when exposed to acute mechanical vortex (a phenotype referred to as bang-sensitivity), though to a lesser extent than following a HS (Figure S2D). For the remaining experiments, we used the dStim RNAi #1 (verified in ^32^, see Methods).

**Figure 4:**
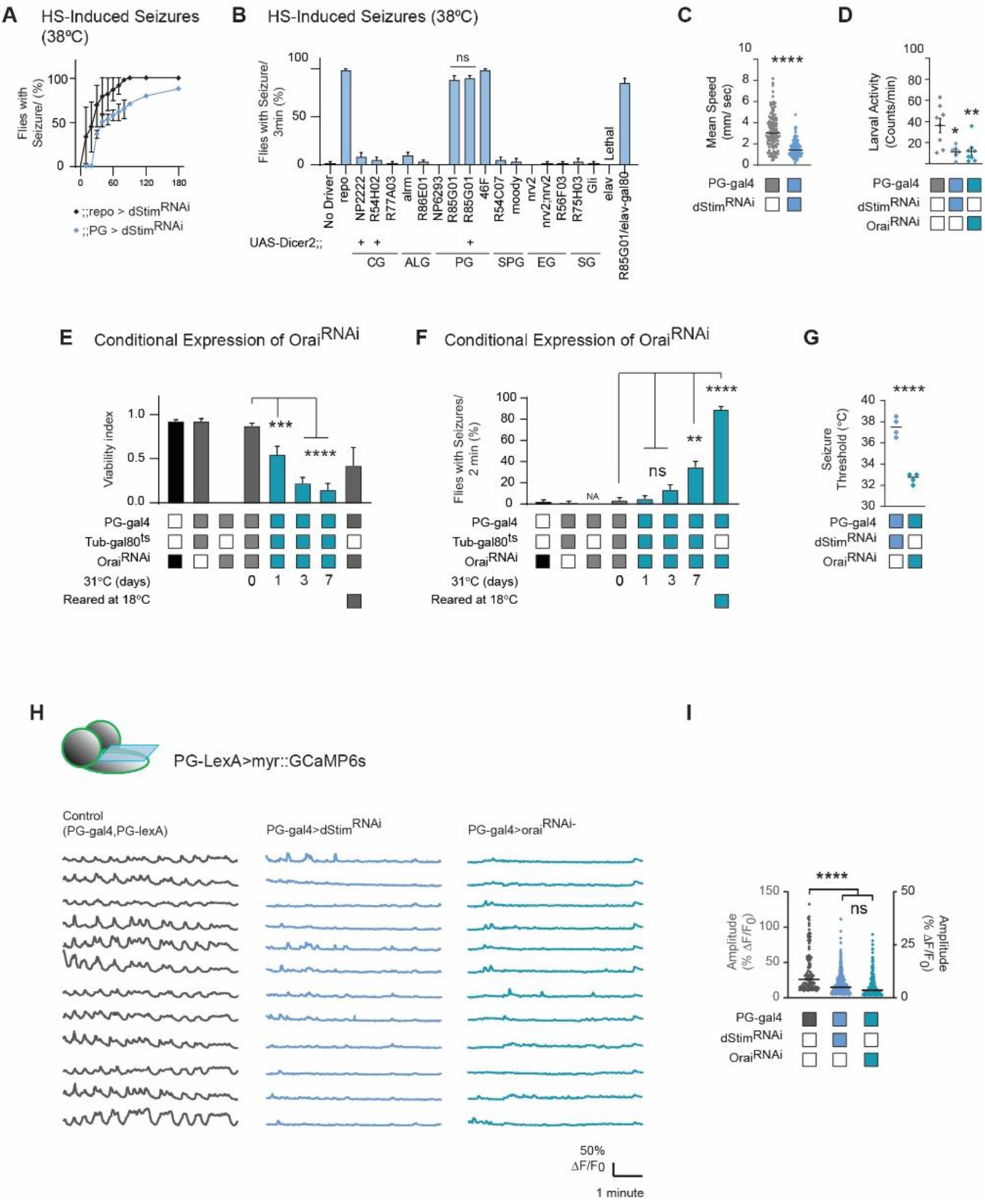
Perineural Knockdown of Store-Operated Ca^2+^ Entry Pathway (SOCE) Components Impairs Locomotor Activity and Increases Seizure Susceptibility. (**A**) Time course of heat-shock induced seizures (38.5°C, HS) for repo>dStim^RNAi^ and PG>dStim^RNAi^ is shown (N=4 groups of 20 flies/genotype, error bars are mean ± SEM).For control, see Video 4. (**B**) Histogram summarizing the percent of flies exhibiting HS-induced seizures at 3 minutes (38.5°C). An array of glial specific gal4 drivers were used to knock down dStim (see methods). Only knockdown of dStim using perineural glia (PG) drivers (46F-gal4 and GMR85G01-gal4) recapitulated the pan-glial HS-induced seizure phenotype. Inhibiting gal4 expression of the RNAi in neurons with gal80 (C155-gal80) does not alter the seizures observed with GMR85G01-gal4 knockdown (N=4 groups of >10 flies/genotype, error bars are mean ± SEM). (**C-D**) Activity level of adults (**C**) and 3^rd^ instar larvae (**D**) expressing the dStim RNAi using the PG driver (N=8 larvae/genotype, N>100 flies/genotype, median is presented). (**E-F**) Perineural conditional knockdown of Orai using gal4/gal80^ts^. Rearing adult flies at the restrictive temperature (>30°C) for gal80^ts^ allows expression of Orai^RNAi^ only in adults (N=4 groups of 20 flies/condition, error bars are mean ± SEM). (**E**) Over the course of several days at 31°C, the majority of PG>Orai^RNAi^/gal80^ts^ flies rapidly deteriorated and died (~80% mortality after 7 days, p<0.0001). (**F**) A significant increase in seizures (p<0.0001) was seen after seven days of rearing flies at the restrictive temperature for gal80^ts^ (31°C). Approximately 30% of surviving adults showed seizures and the rest displayed severe locomotor defects. (**G**) Behavioral analysis of HS-induced seizures revealed that PG>Orai^RNAi^ larvae have a significantly lower temperature threshold (p<0.0001) relative to PG>dStim^RNAi^ animals (N=4 groups of 20 animals/genotype/temperature, median is presented). (**H-I**) Imaging of PG-lexA>LexApo-myr::GCaMP6s following genetic inhibition of SOCE (see Methods). (**H**) Representative traces of mean myr::GCaMP6s fluorescence (% Δ*F/F*_0_) in ROIs assigned to VNC hemi-segments show significant reduction in both amplitude (p<0.0001) and frequency in SOCE knockdowns (PG>dStim^RNAi^ and PG>Orai^RNAi^). (**I**) Histogram summarizing myr::GCaMP6s amplitudes in control (n=106 transients), PG>dStim^RNAi^ (n=190) and PG>Orai^RNAi^ (n=190) VNCs (n=12 ROIs/4 animals). **=p<0.01, ***=p<0.001, ****=p<0.0001, Student’s t-test.

The seizure phenotype observed from pan-glial knockdown of dStim might result from a developmental role for glial dStim in the CNS. To test for a developmental effect of dStim knockdown, we conditionally expressed a single copy of dStim RNAi using gal4/gal80^ts^ (see Methods) only in adult flies. Adult flies reared at the permissive temperature for gal80^ts^ for 3 days (>30°C, to allow dStim RNAi expression) exhibited significantly more seizures, with ~60% of flies displaying seizure phenotypes (Figure S2E). dStim, together with the calcium release-activated calcium channel protein, Orai, are implicated in the SOCE pathway. Pan-glial knockdown of Orai was embryonic lethal, while dStim knockdown was largely viable and showed seizure phenotypes as described above (Figure S2F). These experiments suggest that the SOCE pathway is likely to be essential in *Drosophila* glia, with a role in regulating neuronal excitability and the susceptibility of seizure-like episodes.

### Knockdown of dStim in Perineurial Glial Cells Increases Seizure Susceptibility

The finding that pharmacological inhibition of ER Ca^2+^ stores significantly reduced PG Ca^2+^ signaling (Figure 3A) suggests that PG Ca^2+^ activity relays on internal ER Ca^2+^ stores. Indeed, knockdown of dStim with the PG driver, recapitulated the HS-induced seizure phenotype observed following pan-glial knockdown (Figure 4A, 4B, Video 4, right). Knockdown of dStim with a second perineurial driver (46F-gal4^33^) showed similar seizure phenotype. Seizure characteristics were slightly different between pan-glial and PG specific knockdowns (Figure 4A), with a longer time required for seizure initiation and slower seizure kinetics in dStim PG-knockdown. Co-expressing Dicer-2 with dStim^RNAi^ did not enhance the phenotype (Figure 4B). The weaker phenotype caused with the PG driver suggests that while the primary requirement for dStim in seizure induction is in PG cells, it may also function in other glial subtypes. Alternatively, the PG driver may result in less efficient knockdown of the transcript compared to the pan-glial driver. To test for the requirement of dStim for seizure initiation in other glial subtypes, we performed a secondary screen in which dStim was knocked down specifically in different glial subpopulations. For this screen, we used a series of glial drivers previously described in the field^28^ (Figure 4B, see Methods). Knockdown of dStim in the two glial subpopulations that are best positioned to influence neuronal activity, cortex glia and astrocyte-like glia, failed to recapitulate the phenotype of the pan-glial knockdown (Figure 4B) and showed no apparent effect on viability or motor activity. These results indicate SOCE’s glial role in neuronal excitability is primarily required in PG cells. To test whether PG>dStim^RNAi^ animals display altered behaviors in the absence of a temperature trigger, basal locomotor activity levels were examined. Adult PG>dStim^RNAi^ animals exhibited significant reduction in activity levels (Figure 4C) at room temperature, indicating basal activity is also impaired with reduced dStim function. Other aspects of motor activity such as larval light avoidance (Figure S2G) and adult phototaxis and geotaxis (Figure S2H, I) were not obviously disrupted.

To further characterize the effects of disrupting SOCE, Orai was specifically knocked down in PG cells. Orai knockdown with the PG driver (PG>Orai^RNAi^) was adult lethal with most animals surviving until late pupal stages. Third instar PG>Orai^RNAi^ larvae showed a significant defect in locomotor activity (Figures 4D) and HS-induced seizure-like activity when placed at 38°C (Video 5, left). PG>dStim^RNAi^ larvae showed similar activity impairment as PG>Orai^RNAi^ larvae (Figures 4D). To exclude the possibility that a developmental effect of Orai knockdown in PG cells leads to lethality, we examined brain morphology in larvae co-expressing Orai^RNAi^ and mCD8::GFP and found no significant changes in brain wrapping by PG cells or brain size of 3^rd^ instar larvae (Figure S2J). To examine this possibility further, Orai^RNAi^ was conditionally expressed with gal4/gal80^ts^ (see Methods) only in adult flies. PG>Orai^RNAi^/gal80^ts^ animals reared at 25°C survived to adulthood and showed no HS-induced seizures 1-day post eclosion (Figures 4E, 4F). However, over the course of several days at 31°C, the majority (~80%) of PG>Orai^RNAi^/gal80^ts^ flies showed progressive loss of motor control and death (Figure 4E), with ~35% of the surviving flies displaying seizures after 7 days (Figure 4F). These results indicate Orai function in PG cells is crucial for normal brain function. Consistent with the more severe phenotype of Orai knockdown compared to dStim knockdown with repo-gal4 or PG-gal4, the temperature threshold for seizures in PG>Orai^RNAi^ larvae was significantly lower compared to PG>dStim^RNAi^ (Figure 4G). This raises the possibility that lethality observed in animals with conditional or constant expression of Orai^RNAi^ is due to persistent severe seizures and impaired locomotion at both 25°C and 31°C. To test this hypothesis, we reduced the expression of Orai^RNAi^ by rearing PG>Orai^RNAi^ animals at 18°C, which is below the optimal activation temperature of the UAS/gal4 system^34^. Under these conditions, ~50% of PG>Orai^RNAi^ animals survived to adulthood and showed no HS-induced seizures following eclosion. However, flies that were moved to 25°C after eclosion showed HS-induced seizures one day later (~80%, Figures 4E, 4F, Video 5, right). Over the course of several days at 25°C, the majority (~80%) of PG>Orai^RNAi^/gal80^ts^ flies rapidly deteriorated and died. The stronger effect of Orai knockdown might be due to a stronger suppression of the SOCE pathway, or due to Orai functioning in a dStim-independent manner^35^. However, the similar seizure and locomotion phenotypes in dStim and Orai knockdown suggest the two genes act in the same pathway. PG knockdown of other central components of ER-related Ca^2+^ signaling (Sarco/Endoplasmic reticulum Ca^2+^-ATPase, SERCA, and inositol 1,4,5-trisphosphate receptor, Itpr/ IP_3_R) revealed that while SERCA knockdown was adult lethal, IP_3_R knockdown was viable (Figure S2K) and showed no apparent behavioral phenotype (Figure S2L). To reduce the expression of SERCA^RNAi^, we raised PG>SERCA^RNAi^ animals at 18°C. Under these conditions, only female PG>SERCA^RNAi^ animals survived to adulthood, but they failed to inflate their wings and died within several hours after eclosion.

Knockdown of the SOCE pathway is predicted to impair restoration of ER Ca^2+^ stores and reduce the amplitudes of ER originated Ca^2+^ transients. To test this hypothesis, PG Ca^2+^ transients were imaged using PG-lexA>myr::GCaMP6s in PG knockdowns of SOCE components (PG-gal4>dStim^RNAi^ and PG-gal4>Orai^RNAi^). Indeed, Ca^2+^ signals from VNC-PG and Brain-PG showed significantly reduced amplitudes in both PG>dStim^RNAi^ and PG>Orai^RNAi^ (Video 6, Figure 4H, I, note that SOCE RNAis are plotted on the right Y axes). Together, these data indicate a critical requirement for SOCE in *Drosophila* PG cells, independent of IP_3_R, that is necessary to maintain normal brain function.

### Knockdown of dStim in Perineurial Glial Cells Does Not Affect Blood-Brain-Barrier Integrity

PG cells are thought to influence the development, integrity and function of the blood-brain-barrier (BBB) formed by the SPG layer, as suggested for astrocytes in the mammalian CNS^36^. Ca^2+^ waves within SPG cells were previously reported to control nutrient-dependent reactivation of *Drosophila* neural stem cells and brain growth^25,26^. Perineurial glia also contribute to the deposition of the neural lamella, thus participating in regulating brain shape and stiffness. Alterations in the neural lamella can disrupt brain shape and migration of PG cells^16^. Hence, alteration in PG Ca^2+^ signaling in dStim and Orai knockdowns could lead to seizures secondary to a role for PG ER related Ca^2+^ signaling in controlling brain development. However, this seems unlikely given conditional knockdown of dStim or Orai in adult flies recapitulates the seizure phenotype (Figures S2E and 4F). To test for an effect of glial SOCE on brain growth and PG migration, we co-expressed dStim^RNAi^ or Orai^RNAi^ together with mCD8::GFP specifically in PG cells and monitored brain growth. We found no significant changes between control, PG>dStim^RNAi^ and PG>Orai^RNAi^ animals, both in brain wrapping by PG and in brain size of third instar larvae (Figure S2J). To test whether PG SOCE knockdown compromises the function of the BBB, we incubated PG>dStim^RNAi^ brains with Alexa647-conjugated 10kD dextran and monitored brain penetrance of the dye. In both parental control and PG>dStim^RNAi^ brains, fluorescent dextran remained at the periphery of the brain (Figure S2M), indicating dStim knockdown does not grossly alter permeability of the BBB, while in SPG>Su(H)^RNAi^ (positive control), significant uptake of the dye was observed, indicating dysfunction of the BBB in these animals. Taken together, these results suggest that PG knockdown of SOCE does not affect the gross integrity of the *Drosophila* blood-brain-barrier.

### The Spread of PG Ca^2+^ Waves Through Gap junctions is Crucial for the Prevention of Seizures

Ca^2+^ waves in mammalian astrocytes can be spread via direct communication between adjoining cells through gap junction channels or by release of gliotransmitters that activate neighboring cells via membrane receptors. These two mechanisms are thought to work in parallel to coordinate Ca^2+^ activity between neighboring cells^37^. To examine the mechanism that mediates Ca^2+^ wave spread within the PG cellular sheet surrounding the brain, we first manipulated PG secretion by overexpressing the temperature-sensitive (ts) allele of the *Drosophila* Dynamin homolog, Shibire (Shi^ts^). Conditionally inhibiting endocytosis (and subsequent exocytosis) in PG cells, either acutely (by subjecting PG>Shi^ts^ flies directly to 38°C) or constitutively (by pre-incubating PG>Shi^ts^ flies at the restrictive temperature, 30°C, before a 38°C HS) had no effect on seizure susceptibility of either wildtype or PG>dStim^RNAi^ animals (Figure 5A, see Methods). These results suggest gap junction communication may be the dominant mode of Ca^2+^ wave spread in *Drosophila* PG cells rather than through secretion of exogenous factors.

**Figure 5:**
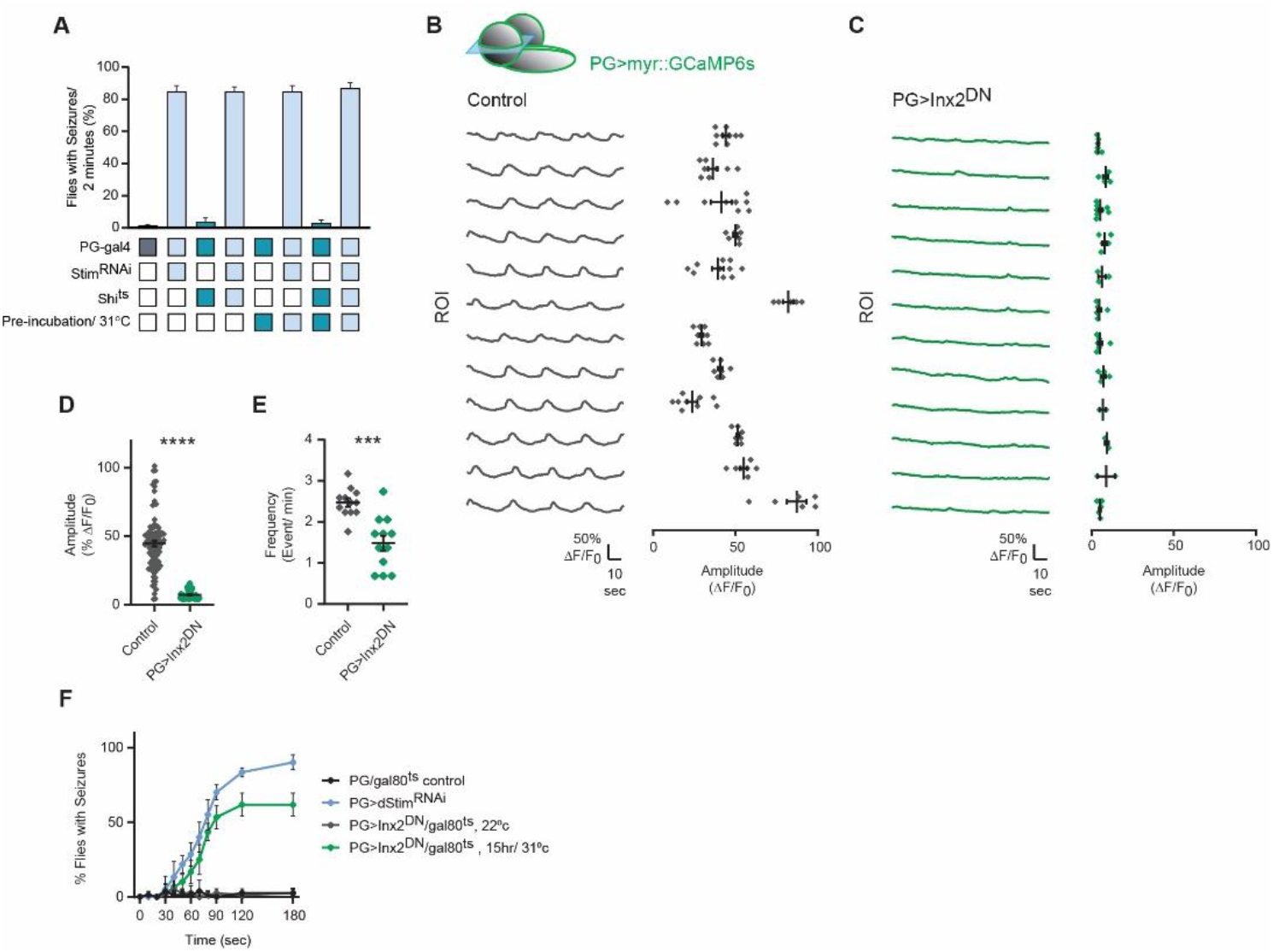
Spread of PG Ca^2+^ Waves Through Gap junctions is Necessary for Preventing Neuronal Hyperexcitability. (**A**) Histogram summarizing the percent of flies showing seizures after a 2-minute HS (38.5°C, HS). PG>Shi^ts^ was used for acute or prolonged inhibition of endo-exocytosis balance. Both conditions had no effect on seizure susceptibility of either wildtype or PG>dStim^RNAi^ flies (N=4 groups of 20 flies/genotype/condition). (**B-E**) Imaging of PG>myr::GCaMP6s in dissected 3^rd^ instar *Drosophila* larvae expressing Inx2^DN^. (**B-C**) Representative traces of mean myr::GCaMP6s fluorescence (% Δ*F/F*_0_) for randomly assigned ROIs show that PG Ca^2+^ waves are abolished in PG>Inx2^DN^ animals compared to controls, although some local activity still remains within individual cells. (**D-E**) Comparisons of amplitude and frequency of Ca^2+^ transients in PG>Inx2^DN^ and controls (n≥60 ROIs/3-5 animals). (**D**) Histogram showing a significant reduction (p<0.0001) in the amplitude of the remaining Ca^2+^ activity in PG>Inx2DN relative to control (n=52 transients for PG>Inx2DN, n=95 transients for control). (**E**) Histogram showing a significant reduction (p<0.001) in the frequency of the remaining Ca^2+^ activity in PG>Inx2^DN^ relative to control (n=20 “active” ROIs/3 animals/genotype). (**F**) Time course of HS-induced seizures (38.5°C, HS) following conditional expression of PG>Inx2^DN^ only in adult flies. Flies in which Inx2^DN^ expression was induced for 15 hours show enhanced seizures relative to controls (N=4 groups of 20 flies/genotype). **=p<0.001, ****=p<0.0001, Student’s t-test.

Within the *Drosophila* BBB, SPG cells exhibit Ca^2+^ waves that spread through neighboring SPG cells via gap junctions^25,26^. Astrocytes and pericytes at the vertebrate BBB also exhibit Ca^2+^ waves that spread through gap junctions^38^. To test whether the spread of Ca^2+^ waves between neighboring PG cells is mediated by gap junctions, we over-expressed a dominant negative (DN) form of one of the *Drosophila* gap junction homologs, Inx2 (Inx2^DN^), in PG cells. PG overexpression of Inx2^DN^ significantly inhibited the spread of Ca^2+^ waves in PG cells at the VNC and within the brain (Video 7), while also significantly reducing both the amplitude and frequency of Ca^2+^ signals within individual PG cells (Figure 5B-E). Similar to other genetic manipulations described above, RNAi-mediated knockdown of either Inx1 or Inx2, or overexpression of Inx2^DN^, were found to be adult lethal. Conditionally expressing Inx2^DN^ with gal4/gal80^ts^ (see Methods) only in adult flies significantly increased their seizure susceptibility, with ~60% of the flies showing seizures after 15 hours at the restrictive temperature for gal80^ts^ (>30°C, Figure 5E, Video 7). Together, these data demonstrate that Ca^2+^ waves spread through neighboring PG cells via gap junctions. Blocking the spread of these Ca^2+^ waves recapitulates the behavioral phenotypes of SOCE knockdown, suggesting propagation of PG Ca^2+^ waves is crucial for maintaining normal brain excitability and preventing seizure activity.

## Discussion

In this study we performed a detailed characterization of *Drosophila* PG Ca^2+^ signaling that reveals PG cells exhibit robust, complex, and dynamic Ca^2+^ activity that varies depending upon which brain territory they occupy. These Ca^2+^ transients are independent of extracellular Ca^2+^ and Tg-sensitive, suggesting they originate from intracellular Ca^2+^ stores rather than from plasma membrane influx as observed in *Drosophila* astrocytes and cortex glia. We find that manipulating ER-related Ca^2+^ signaling by disrupting store-operated Ca^2+^ entry (SOCE) or inhibiting the spread of Ca^2+^ waves through gap junctions significantly reduces PG Ca^2+^ activity. Ca^2+^ signaling through this pathway in PG cells is essential for controlling neuronal excitability, as knockdown of SOCE components or manipulation of gap junction function impairs basal motor activity and increases seizure susceptibility. These data indicate PG Ca^2+^ signaling involves store dependent Ca^2+^ signaling and is essential for maintaining normal nervous system function. As such, glial Ca^2+^-dependent signaling through either microdomain extracellular Ca^2+^ influx or Ca^2+^ release from internal stores are both essential for regulating neuronal function depending upon the glial subtype.

A key question moving forward is how PG Ca^2+^ waves mechanistically regulate brain function. Maintenance of neuronal excitability requires a fine-tuned extracellular ion balance and a steady supply of nutrients and metabolites. This homeostasis is achieved by evolutionary conserved specialized structures that form the blood-brain barrier (BBB). The primary function of the BBB is to maintain homeostasis by regulating influx and efflux transport, a role that requires tight cell-to-cell interactions. The mammalian BBB consists of endothelial cells, astrocytes, pericytes, neurons, and microglia which shape the homeostatic function of the barrier^12^. Several components of the BBB, including endothelial cells, astrocytes and pericytes exhibit fluctuations in intracellular Ca^2+^, suggesting a generalized role for Ca^2+^ activity in BBB function^37,39–41.^ In *Drosophila*, the BBB is formed by two glial layers: the PG and SPG cells (Figure 1A). The main barrier function is attributed to SPG cells that form pleated septate junctions and prevent paracellular diffusion, similar to tight junctions in the mammalian endothelial BBB. PG establish the first diffusion barrier and provide structural roles (i.e. secretion of the neural lamella and providing rigidity to the CNS) and provide SPG with metabolic support, although their exact contribution to BBB function is not fully understood^15,16,20,42.^ We did not find defects in gross morphology of PG or SPG cells when PG Ca^2+^ waves were disrupted. Similarly, the BBB diffusion barrier as assayed with dextran dye penetration was unaffected. As such, PG Ca^2+^ waves are likely to regulate either small molecule diffusion, SPG cell function or secretion of unknown factors from PG cells to shape neuronal excitability.

While interference with SOCE and spread of Ca^2+^ waves across the PG sheet alters behavior and increases seizure susceptibility, the mechanism(s) downstream of intracellular Ca^2+^ changes that alter neuronal excitability is unknown. Based on our current observations, the different signatures in PG Ca^2+^ signaling across different brain regions suggest that although all PG cells utilize ER-store-dependent Ca^2+^ signaling, distinct PG cell populations may employ intracellular Ca^2+^ signaling in unique ways. The *Drosophila* hemolymph-brain barrier (PG and SPG) expresses numerous transporters and receptors that selectively move nutrients, metabolites and other compounds in and out of the brain^13^. Hence, one possible mechanism by which PG Ca^2+^ could alter neuronal activity is through regulation of transport across the BBB. First, PG Ca^2+^ activity could modulate exocytotic/endocytotic cycling of membrane proteins within the PG layer itself, similar to its role in regulating surface levels of K^+^ channels in cortex glia^24^ and GABA transporters in astrocyte-like glia^22^. Second, PG Ca^2+^ may regulate transport in SPG cells via a cell non-autonomous mechanism. A recent study found that efflux transporters in *Drosophila* SPG cells were regulated by a circadian clock in PG cells ^43^. Though the gross anatomy of the BBB and basic diffusion of large molecules across the barrier were not affected, regulation of more subtle aspects of BBB transport of small molecules such as ions and metabolites could be altered. PG cells also play structural roles by secreting proteins composing the neural lamella and providing rigidity to the CNS^16^. As such, impairments in the fine structure of the BBB might indirectly affect transport. At the *Drosophila* neuromuscular junction (NMJ), Ca^2+^ release from the ER is involved in microtubule stabilization^44^, and disruption in this process impairs synaptic growth, synaptic vesicle release probability and decrease synaptic transmission. Thus, loss of PG SOCE and subsequent depletion of ER Ca^2+^ stores might directly influence the secretion of proteins that compose the neural lamina, or induce destabilization of PG microtubules and alter rigidity that is crucial for brain homeostasis and function^16^. Further studies will be required to define how PG Ca^2+^ waves ultimately control neuronal excitability and whether different PG populations use distinct intracellular Ca^2+^ signaling pathways that are dependent on their unique Ca^2+^ wave dynamics.

Single-cell transcriptomic analyses of mammalian glial subtypes, including astrocytes, have advanced our understanding of astrocyte diversity. Astrocytes from different brain regions, as well as within the same region, have distinct transcriptomic profiles that allow classification into novel subpopulations with unique spatial distribution and signaling pathways^2,45^. To date, no molecular differences have been described between PG cells derived from the CNS or PNS, and despite morphological differences, PG cells are thought to share similar functional properties^16^. Transcriptomic analysis of *Drosophila* surface glia (PG and SPG together) demonstrated these cells collectively show molecular signatures similar to vertebrate brain-vascular endothelial cells that form the BBB^13^. However, this analysis lacked single-cell resolution required for molecular distinction of PG cells from different brain areas. Single cell transcriptomic analysis of the *Drosophila* brain displayed relatively low coverage of PG cells (~70 cells,^46^), preventing any distinction between possible subpopulations. Our data suggest PG cells derived from different brain regions can be distinguished based on Ca^2+^ wave dynamics, and further transcriptomic analyses might yield insights into the diversity of PG cells and glial cells in general.

Although astrocytes were traditionally considered to serve only supportive functions in the brain, the discovery of astrocytic intracellular Ca^2+^ signals has changed our view of how these cells contribute to brain function. Accumulating data indicate astrocytes can respond to neuronal activity and regulate neuronal function via intracellular astrocytic Ca^2+^ signaling. However, the functional consequences of glial Ca^2+^ signaling on neuronal physiology and brain function are not fully understood. One of the controversies in the field of glial biology is the functional distinction between ER-mediated somatic Ca^2+^ oscillations and near-membrane microdomain Ca^2+^ oscillations in glial processes. A central mechanism in intracellular Ca^2^ signaling is the SOCE pathway which re-fills ER Ca^2+^ stores upon depletion triggered by a signaling cascade. In this pathway, the gating of the plasma membrane Ca^2+^ channel, Orai, is controlled by the ER localized Ca^2+^ sensor, Stim, leading to Ca^2+^ influx and restoration of the ER Ca^2+^ store. While Orai and Stim expression have been detected in mammalian astroglia, the role of SOCE in glial biology is yet to be fully characterized. Our functional analysis indicate SOCE is a critical Ca^2+^ signaling pathway in PG cells that can act independently of IP3 receptors.

Accumulating evidence indicate glia are likely to play a central role in a host of neurological disorders, with multiple disease-associated genes enriched in glial subtypes^47^. Studies investigating the mechanisms underlying epileptic seizures have primarily focused on neuronal origins, though accumulating evidence highlights an important role of non-neuronal cells in both the generation and spread of epileptic seizures in the brain. In this study, we found that alterations in the SOCE pathway in the *Drosophila* BBB lead to seizures-like episodes (referred to as “seizures”) without affecting basic barrier function. This suggests that SOCE within the BBB regulates more subtle processes mediated by the barrier, such of the regulation of active transport of small molecules or ions. In vertebrates, a tight connection between seizures and BBB dysfunction has also been found, with some studies showing that prolonged seizures or brain injury can lead to changes in BBB properties and subsequent BBB dysfunction, and other studies suggesting a causative role for BBB dysfunction in epileptogenesis^48,49^. Future characterization of how glial Ca^2+^ signaling within the *Drosophila* BBB actively shapes neuronal excitability should shed light on the broader role of BBB function in the generation of seizures and suggest potential new treatment targets for epilepsy.

## Methods

### Key resources table

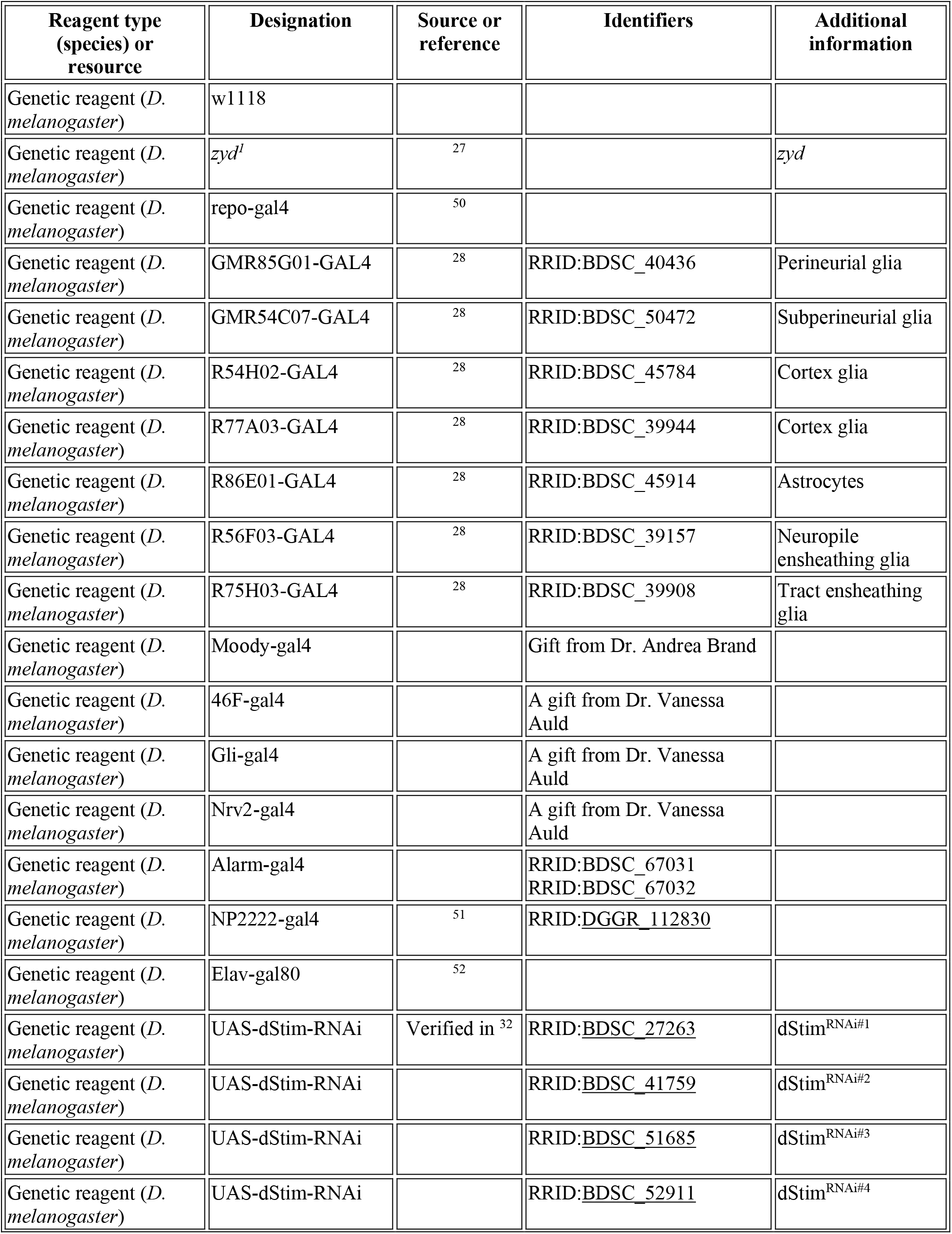
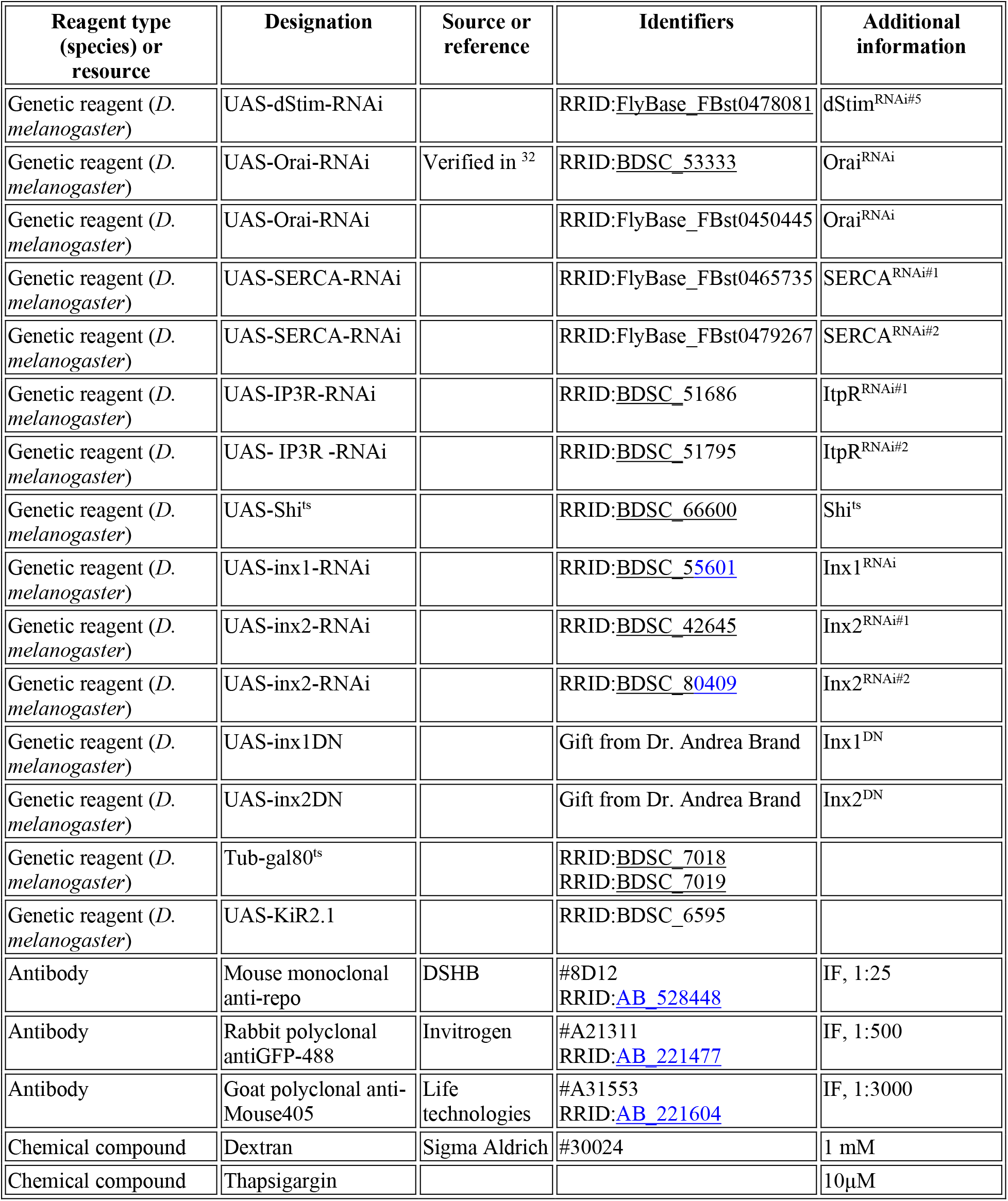

| Reagent type (species) or resource | Designation | Source or reference | Identifiers | Additional information |
| --- | --- | --- | --- | --- |
| Genetic reagent (*D. melanogaster*) | w1118 |  |  |  |
| Genetic reagent (*D. melanogaster*) | *zyd<sup>l</sup>* | 27 |  | *zyd* |
| Genetic reagent (*D. melanogaster*) | repo-gal4 | 50 |  |  |
| Genetic reagent (*D. melanogaster*) | GMR85G01-GAL4 | 28 | RRID:BDSC_40436 | Perineurial glia |
| Genetic reagent (*D. melanogaster*) | GMR54C07-GAL4 | 28 | RRID:BDSC_50472 | Subperineurial glia |
| Genetic reagent (*D. melanogaster*) | R54H02-GAL4 | 28 | RRID:BDSC_45784 | Cortex glia |
| Genetic reagent (*D. melanogaster*) | R77A03-GAL4 | 28 | RRID:BDSC_39944 | Cortex glia |
| Genetic reagent (*D. melanogaster*) | R86E01-GAL4 | 28 | RRID:BDSC_45914 | Astrocytes |
| Genetic reagent (*D. melanogaster*) | R56F03-GAL4 | 28 | RRID:BDSC_39157 | Neuropile ensheathing glia |
| Genetic reagent (*D. melanogaster*) | R75H03-GAL4 | 28 | RRID:BDSC_39908 | Tract ensheathing glia |
| Genetic reagent (*D. melanogaster*) | Moody-gal4 |  | Gift from Dr. Andrea Brand |  |
| Genetic reagent (*D. melanogaster*) | 46F-gal4 |  | A gift from Dr. Vanessa Auld |  |
| Genetic reagent (*D. melanogaster*) | Gli-gal4 |  | A gift from Dr. Vanessa Auld |  |
| Genetic reagent (*D. melanogaster*) | Nrv2-gal4 |  | A gift from Dr. Vanessa Auld |  |
| Genetic reagent (*D. melanogaster*) | Alarm-gal4 |  | RRID:BDSC_67031<br>RRID:BDSC_67032 |  |
| Genetic reagent (*D. melanogaster*) | NP2222-gal4 | 51 | RRID:DGGR_112830 |  |
| Genetic reagent (*D. melanogaster*) | Elav-gal80 | 52 |  |  |
| Genetic reagent (*D. melanogaster*) | UAS-dStim-RNAi | Verified in <sup>32</sup> | RRID:BDSC_27263 | dStim<sup>RNAi#1</sup> |
| Genetic reagent (*D. melanogaster*) | UAS-dStim-RNAi |  | RRID:BDSC_41759 | dStim<sup>RNAi#2</sup> |
| Genetic reagent (*D. melanogaster*) | UAS-dStim-RNAi |  | RRID:BDSC_51685 | dStim<sup>RNAi#3</sup> |
| Genetic reagent (*D. melanogaster*) | UAS-dStim-RNAi |  | RRID:BDSC_52911 | dStim<sup>RNAi#4</sup> |
| Genetic reagent (*D. melanogaster*) | UAS-dStim-RNAi |  | RRID:FlyBase_FBst0478081 | dStim<sup>RNAi#5</sup> |
| Genetic reagent (*D. melanogaster*) | UAS-Orai-RNAi | Verified in <sup>32</sup> | RRID:BDSC_53333 | Orai<sup>RNAi</sup> |
| Genetic reagent (*D. melanogaster*) | UAS-Orai-RNAi |  | RRID:FlyBase_FBst0450445 | Orai<sup>RNAi</sup> |
| Genetic reagent (*D. melanogaster*) | UAS-SERCA-RNAi |  | RRID:FlyBase_FBst0465735 | SERCA<sup>RNAi#1</sup> |
| Genetic reagent (*D. melanogaster*) | UAS-SERCA-RNAi |  | RRID:FlyBase_FBst0479267 | SERCA<sup>RNAi#2</sup> |
| Genetic reagent (*D. melanogaster*) | UAS-IP3R-RNAi |  | RRID:BDSC_51686 | ItpR<sup>RNAi#1</sup> |
| Genetic reagent (*D. melanogaster*) | UAS- IP3R -RNAi |  | RRID:BDSC_51795 | ItpR<sup>RNAi#2</sup> |
| Genetic reagent (*D. melanogaster*) | UAS-Shi<sup>ts</sup> |  | RRID:BDSC_66600 | Shi<sup>ts</sup> |
| Genetic reagent (*D. melanogaster*) | UAS-inx1-RNAi |  | RRID:BDSC_55601 | Inx1<sup>RNAi</sup> |
| Genetic reagent (*D. melanogaster*) | UAS-inx2-RNAi |  | RRID:BDSC_42645 | Inx2<sup>RNAi#1</sup> |
| Genetic reagent (*D. melanogaster*) | UAS-inx2-RNAi |  | RRID:BDSC_80409 | Inx2<sup>RNAi#2</sup> |
| Genetic reagent (*D. melanogaster*) | UAS-inx1DN |  | Gift from Dr. Andrea Brand | Inx1<sup>DN</sup> |
| Genetic reagent (*D. melanogaster*) | UAS-inx2DN |  | Gift from Dr. Andrea Brand | Inx2<sup>DN</sup> |
| Genetic reagent (*D. melanogaster*) | Tub-gal80<sup>ts</sup> |  | RRID:BDSC_7018<br>RRID:BDSC_7019 |  |
| Genetic reagent (*D. melanogaster*) | UAS-Kir2.1 |  | RRID:BDSC_6595 |  |
| Antibody | Mouse monoclonal anti-repo | DSHB | #8D12<br>RRID:AB_528448 | IF, 1:25 |
| Antibody | Rabbit polyclonal antiGFP-488 | Invitrogen | #A21311<br>RRID:AB_221477 | IF, 1:500 |
| Antibody | Goat polyclonal anti-Mouse405 | Life technologies | #A31553<br>RRID:AB_221604 | IF, 1:3000 |
| Chemical compound | Dextran | Sigma Aldrich | #30024 | 1 mM |
| Chemical compound | Thapsigargin |  |  | 10μM |

### *Drosophila* Genetics and Molecular Biology

Flies were cultured on standard medium at 22°C unless otherwise noted. The UAS/gal4 and LexAop/LexA systems were used to drive transgenes in glia using the indicated drivers. The *UAS-dsRNAi* flies used in the study were obtained from the VDRC (Vienna, Austria) or the TRiP collection (Bloomington *Drosophila* Stock Center, Indiana University, Bloomington, IN, USA). UAS-*myrGCaMP6s* was constructed by replacing GCaMP5 in the previously described myrGCaMP5 transgenic construct ^27^. To generate UAS- and lexAop-OER:GCaMP6f flies, OER:GCaMP6f (gift from Mikoshiba Hiroko, ^31^) was subcloned into either pBID-UASc or pBID-LexAop plasmids using standard methods (Epoch Life Science Inc.). Transgenic flies were obtained by standard germline injection (BestGene Inc). For all experiments described, both male and female larvae or adults were used, unless otherwise noted. For survival assays, embryos were collected in groups of ~50 and transferred to fresh vials (n=3). 3^rd^ instar larvae and/or pupae were counted. Survival rate (SR) was calculated as:

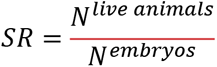

For conditional expression using Tub-gal80^ts^ (Figures 4E-F, S2E and 5E), animals of the designated genotype were reared at 22°C with gal80 suppressing gal4-driven transgene expression (dStim^RNAi^, Orai^RNAi^ and inx2^DN^). Adult flies were then transferred to a 31°C incubator to inactivate gal80 and allow gal4 knockdown for the indicated period. For UAS/gal4 inhibition by low temperature (Figures 4E, F), PG> Orai^RNAi^ animals were reared at 18°C to suppress gal4-driven transgene expression. Adult flies were transferred to a 25°C incubator upon eclosion to allow gal4 knockdown/overexpression for the indicated period. For inhibiting transgene expression specifically in neurons (Figure 2A), elav-gal80^52^ was used.

### Behavioral analysis

For assaying temperature-sensitive seizures, adult males aged 1-2 days were transferred in groups of ~10-20 flies (n ≥ 3, total # of flies tested in all assays was always >40) into preheated vials in a water bath held at the indicated temperature with a precision of 0.1°C. Seizures were defined as the condition in which the animal lies incapacitated on its back or side with legs and wings contracting vigorously^27^. For screening purposes, only flies that showed normal wildtype-like behavior (i.e. walking up and down on vial walls) after >2min of heat-shock were counted as not seizing. For assaying seizures in larvae, 3^rd^ instar larvae were gently washed with PBS and transferred to 1% agarose plates or empty fly vials and heated to 38°C. Larval seizures were defined as continuous unpatterned contraction of the body wall muscles that prevented normal crawling behavior ^27^. For determining seizure temperature threshold, groups of 10 animals were heat-shocked to the indicated temperature (ranging 30-39.0°C in 0.5°C increments). Threshold was defined as the temperature in which > 50% of the animals were seizing after 1 minute.

For assaying bang sensitivity, adult male flies in groups of ~10-20 (n=3) were assayed 1-2 days post-eclosion. Flies were transferred into empty vials and allowed to rest for 1-2 hr. Vials were vortexed at maximum speed for 10 seconds and the number of flies that were upright and mobile was counted at 10 s intervals.

For larval light avoidance, assays were performed using protocols previously described with minor modifications. Briefly, pools of ~20 3^rd^ instar larvae (108-120 hr after egg laying) were allowed to move freely for 5 minutes on Petri dishes with settings for the phototaxis assay. Petri dish lids were divided into quadrants with two quadrants blackened to create a dark environment. The number of larvae in light versus dark quadrants was then scored (n=4). Response indices (RI) were calculated as:

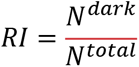

For adult positive phototaxis, assays were performed using protocols described previously following minor modifications. Briefly, pools of ~20 flies (3-5 days after eclosion) were allowed to move freely for 5 minutes on the set up described above for larval light avoidance. The number of flies in light versus dark quadrants was then scored (n=4). Response indices (RI) were calculated as:

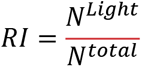

For adult negative geotaxis assays, vials containing pools of ~20 flies (3-5 days after eclosion) were gently tapped and flies were allowed to climb upward for 10 seconds. The number of larvae in the top third of the vial was then scored (n=4). Response indices (RI) were calculated as:

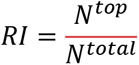

For activity monitoring, wandering 3^rd^ instar larval activity was assayed using a multi-beam system (MB5, TriKinetics) as previously described ^53^. Briefly, individual animals were inserted into 5 mm × 80 mm glass pyrex tubes. Activity was recorded following a 5 minutes acclimation period. Throughout each experiment, animals were housed in a temperature- and light-controlled incubator (25°C, ~40–60% humidity). Post-acquisition activity analysis was performed using Excel to calculate activity level across 1-minute time bins. Each experimental run contained eight control animals and eight experimental animals with n ≥ 3.

### Electrophysiology

Intracellular recordings of wandering 3^rd^ instar male larvae were performed in HL3.1 saline (in mm: 70 NaCl, 5 KCl, 4 MgCl_2_, 1.5 CaCl_2_, 10 NaHCO_3_, 5 Trehalose, 115 sucrose, 5 HEPES-NaOH, pH 7.2) using an Axoclamp 2B amplifier (Molecular Devices) at muscle fiber 6/7 of segments A3-A5. For recording CPG output, the CNS and motor neurons were left intact. Temperature was controlled with a Peltier heating device and continually monitored with a microprobe thermometer.

### In vivo Ca^2+^ imaging

myr::GCaMP6s and ER::GCaMP6f were expressed in PG cells using the drivers described above. PG-gal4 and UAS-myrGCaMP6s was used for most experiments, except for imaging in SOCE knockdowns (PG>dStim^RNAi^ and PG>Orai^RNAi^, Figure 4H, I) where PG-lexA and LexAop-myr::GCaMP6s was used. For live imaging of undissected 2^nd^ instar larvae, animals were washed with PBS and placed on a glass slide with a small amount of Halocarbon oil #700 (LabScientific). Larvae were turned ventral side up and gently pressed with a coverslip and a small iron ring to inhibit movement. For imaging of semi-dissected brains, 3^rd^ instar larvae were dissected in HL3.1 saline (in mm: 70 NaCl, 5 KCl, 4 MgCl_2_, 0.2 CaCl_2_, 10 NaHCO_3_, 5 Trehalose, 115 sucrose, 5 HEPES-NaOH, pH 7.2). A small incision was made above the brain, with the rest of the organs left largely intact. Images were acquired with a PerkinElmer Ultraview Vox spinning disk confocal microscope and a high-speed EM CCD camera at 8-12 Hz with a 20× water-immersion objective using Volocity Software. Single optical planes on the surface or a mid-section of the ventral nerve cord (VNC) or brain hemisphere were imaged. Average myrGCaMP6s signal in perineural glia was quantified in the central thoracic and abdominal segments of the VNC and brain with manually selected regions of interest (ROIs). Ca^2+^ oscillations were analyzed within the first 3 minutes of imaging at room temperature and then normalized to the ROI area.

### Blood-Brain-Barrier Permeability Assay

3^rd^ instar larval brains were dissected in HL3.1 and incubated with Alexa fluor 647-conjugated 10 Kd dextran (Sigma Aldrich #30024) for 5 minutes before image acquisition. Brains were then fixed in 4% PFA in PBS for 5 minutes, washed briefly in PBS, mounted in VectaShield H-1000 (Vector Laboratories) and imaged by confocal microscopy. Subperineural knockdown of Su(H) was used as positive control in each batch.

### Statistical analysis

No statistical methods were used to predetermine sample size. All *n* numbers represent biological replicates. Data were pooled from 2 to 3 independent experiments. Ca^2+^ imaging experiments were randomized and blinded. Students’ t-test was used and P values are represented as *=P<0.05, **=P<0.01, ***=P<0.001, ****=P<0.0001. P<0.05 was considered significant. Data are expressed as mean±SEM or the median.

## Acknowledgements

This work was supported by NIH grants NS40296 and MH104536 to JTL and the Shamir Fellowship form the Israeli Ministry for Science and Technology to SW. We thank the Bloomington *Drosophila* Stock Center (NIH P40OD018537), the Vienna *Drosophila* RNAi Center, the Harvard TriP Project, Vanessa Auld (University of British Columbia), Andrea Brand (Gurdon Institute) for providing *Drosophila* strains, Mikoshiba Hiroko for providing OER:GCaMP6f contract, and members of the Littleton and Parnas labs for helpful discussions and comments on the manuscript.

## Supplementary Information

**Figure supplement 1:**
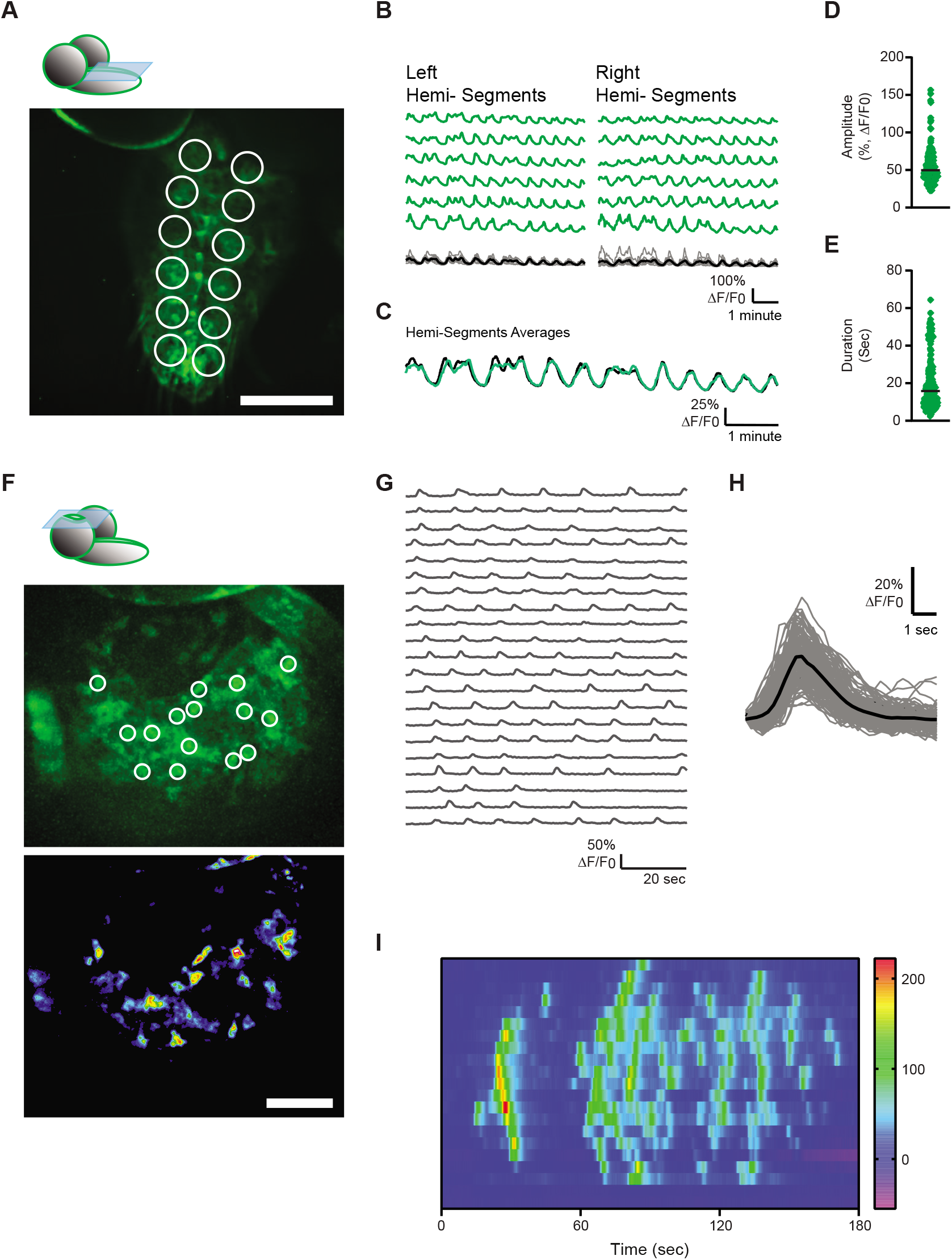
Characterization of PG Ca^2+^ in Cells that Occupy Different CNS Territories. Imaging of PG>myr::GCaMP6s in dissected 3^rd^ instar wildtype *Drosophila* larvae. (**A**) Representative ROI assignment of hemi-segments at the dorsal surface of the VNC. Scale bar, 100 μm. Schematic representation of the *Drosophila* larval brain (top) shows the relative field of view at the dorsal surface of the VNC (light blue). (**B**) Representative traces of mean myr::GCaMP6s fluorescence (% Δ*F/F*_0_) of single hemi-segments reveal dynamic and complex Ca^2+^ activity that is synchronized between hemi-segments. Bottom traces show superimposition of all traces within the same hemi-VNC (n=6, grey) with the mean (black) and (**C**) superimposition of the means of two parallel hemi-VNCs. (**D-E**) Summary of the amplitude (**D**) and duration (**E**) of Ca^2+^ transients at VNC hemi-segments (n=216 transients/48 ROIs/4 animals). (**F**) Representative ROI assignment of single cells at the dorsal surface of a brain hemisphere. ROIs were assigned based on activity map (top). Scale bar, 40 μm. Schematic representation of the *Drosophila* larval brain (top) shows the relative field of view at the surface of a brain hemisphere (light blue). (**G**) Representative traces of mean myr::GCaMP6s fluorescence (% Δ*F/F*_0_) of ROIs shown in (F). (**H**) Superimposition of single brain-PG Ca^2+^ transients (gray) and their mean (black; n=155 transients/50 ROIs/3 animals). (**I**) Heat map representation of the same data as in Figure 2F shows spread of Ca^2+^ waves through adjacent brain-PG areas.

**Figure supplement 2:**
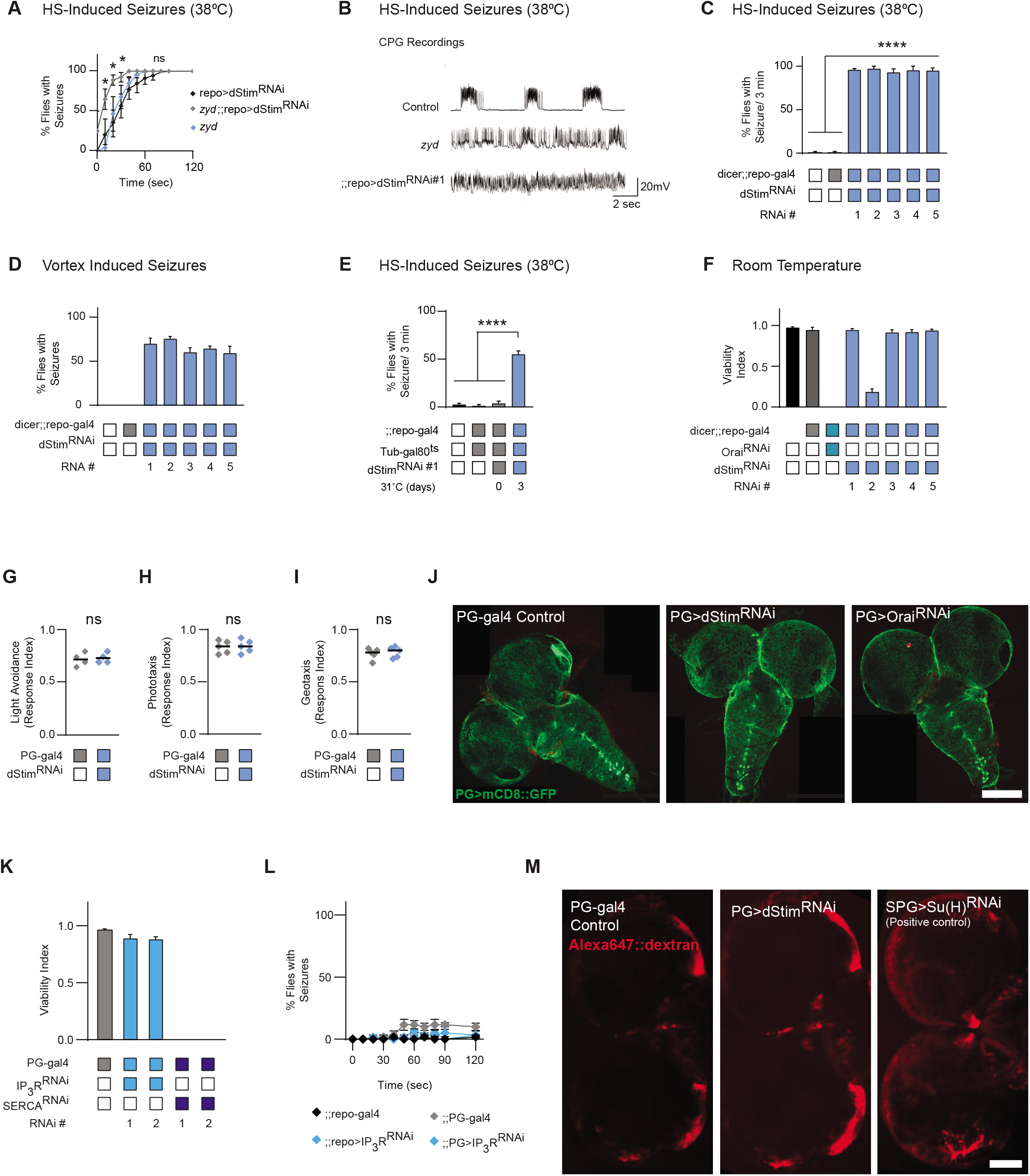
Pan-glial Knockdown of dStim Increases Stress-Induced Seizures. (**A**) Time course of HS-induced seizures (38.5°C, HS) following chronic knockdown of dStim using a pan-glial driver (repo-gal4). Adults containing the *zyd* mutation (*zyd*;;repo>dStim^RNAi^) show enhanced seizures relative to repo>dStim^RNAi^ adults alone (p<0.05, N=4 groups of 20 flies/genotype). (**B**) Representative voltage traces of spontaneous CPG activity recorded at larval 3^rd^ instar muscle 6 at 38°C in wildtype, *zyd* and *zyd*;;repo>dStim^RNAi^ animals (n≥5 preparations/genotype). (**C**) Pan-glial knockdown of dStim with 5 non-overlapping RNAi hairpins (p<0.0001, see Methods, N=4 groups of >10 flies/genotype). (**D**) Behavioral analysis of vortex-induced seizures at 10 sec after stimulus termination (N=3 groups of 20 flies/genotype). (**E**) Histogram summarizing the percent of flies exhibiting heat-shock induced seizures at 3 minutes (38.5°C) following conditional pan-glial knockdown of dStim. Rearing adult flies at the restrictive temperature (>30°C) with gal80^ts^ allows expression of dStim^RNAi^ only at the adult stage. These manipulations partially reproduce the repo>dStim^RNAi^ seizure phenotype (p<0.0001, N=4 groups of 20 flies/genotype). (**F**) Viability analysis of the different RNAi lines used in this study. RNAi was expressed using a pan-glial (repo-gal4) driver (p<0.0001 for dStim^RNAi#2^ and Orai^RNAi^, N=3 groups of 50 embryos/genotype). (**G**) Light avoidance response of control and PG>dStim^RNAi^ 3^rd^ instar larvae (N=4 groups of 20 larvae/genotype). (**H**) Positive phototaxis response of control and PG>dStim^RNAi^ adult flies (N=4 groups of 20 larvae/genotype). (**I**) Negative geotaxis response of control and PG>dStim^RNAi^ adult flies (N=4 groups of 20 larvae/genotype). (**J**) Immunofluorescence imaging reveals no apparent morphological changes in perineural glial wrapping of the CNS or in brain size of 3^rd^ instar larvae (green: anti-GFP, mCD8:GFP, perineural glial membrane). Scale bar=100 μm. (**K**) Viability analysis of SERCA and ItpR/IP_3_R driven with PG-gal4 (N=3 groups of 50 embryos/genotype, error bars are mean±SEM). (**L**) HS-induced seizure assays (38.5°C, HS) for repo>ItpR^RNAi^ and PG>ItpR^RNAi^ are shown (N=4 groups of 20 flies/genotype, error bars are mean±SEM). (**M**) Permeability assay of the BBB measured by immunofluorescent 10 kDa dextran uptake in dissected 3^rd^ instar larval brains reveals no impairment in brain permeability of PG>dStimRNAi brains relative to controls. Note the enhanced red signal in the brains of SPG>Su(H)^RNAi^ larvae, which have a disrupted BBB and served as a positive control. Scale bar=100 μm. *=p<0.05, **=p<0.01, ***=p<0.001, ****=p<0.0001, Student’s t-test.

## Video Files

**Video 1:** (**A**) Representative movie showing spontaneous GCaMP6s events (green) in a live, semi-dissected 3^rd^ instar larval VNC-PG (dorsal surface). Scale bar, 50μm. Video is 4x real time. (**B, C**) Representative movie showing spontaneous GCaMP6s events (green) in a live, semi-dissected 3^rd^ instar larval VNC-PG (dorsal surface). Single cells are mark with nucleus-localized mCherry (mCherry.nls, magenta). Scale bar, 20μm. Video is 4x real time.

**Video 2:** (**A**) Representative movie showing spontaneous GCaMP6s events (green) in PG cells on the surface of the brain of a live, semi-dissected 3^rd^ instar larva (dorsal surface). Scale bar, 50μm. Video is 3x real time. (**B**) Representative movie showing spontaneous PG GCaMP6s events and spread of waves across cells (green) in a section through a brain hemisphere of a live, semi-dissected 3^rd^ instar larva. Scale bar, 50μm. Video is 5x real time. (**C**) Representative movie showing spontaneous PG GCaMP6s events and spread of waves (green) in a section through brain hemisphere of a live, semi-dissected 3^rd^ instar larva treated with 10μM Thapsigargin (Tg). Scale bar, 50μm. Video is 5x real time.

**Video 3:** (**A**) Representative movie showing spontaneous ER::GCaMP6s in a single VNC-PG cell (ventral surface) in live, non-dissected 2^nd^ instar larva. Scale bar, 50μm. Video is 4x real time. (**B**) Representative movie showing spontaneous ER::GCaMP6s events and spread of waves in a PG cell that enwrap the brain in live, non-dissected 3^rd^ instar larva (left). The inset shows the spread of Ca^2+^ related activity between three neighboring cells (right). Scale bars, 50μm (left), 10μm (right). Videos are 4x real time.

**Video 4:** The response of wildtype (Left), pan-glial (repo-gal4, middle) dStim knockdown and perineurial-glial (PG-gal4, right) dStim knockdown adults to a 38.5°C HS is shown. Video is 4x real time.

**Video 5:** (**A**) The response of perineural-glial (PG-gal4) Orai knockdown 3^rd^ instar larvae to a 38.5°C HS is shown. Video is 2x real time. (**B**) The response of perineural-glial (PG-gal4) Orai knockdown adults to a 38.5°C HS is shown. RNAi expression was conditionally induced only in adult flies for 3 days prior to video acquisition. Video is 4x real time.

**Video 6**: (**A**) Representative movie showing spontaneous PG GCaMP6s events are abolished in the brain of a live, semi-dissected 3^rd^ instar PG>Orai^RNAi^ larva. Scale bar, 50μm. Video is 4x real time. (**B**) Representative movie showing spontaneous PG GCaMP6s events are significantly reduced in the VNC of a live, semi-dissected 3^rd^ instar PG>dStim^RNAi^ larva. Scale bar, 50μm. Video is 4x real time.

**Video 7:** (**A**) Representative movie showing spontaneous PG GCaMP6s events are abolished in the brain of a live, semi-dissected 3^rd^ instar PG>Inx2^DN^ larva. Scale bar, 50μm. Video is 4x real time. (**B**) The response of flies conditionally expressing Inx2^DN^ in perineural-glial (PG-gal4) to a 38.5°C HS is shown. Inx2^DN^ expression was conditionally induced only in adult flies for 15 hours prior to video acquisition. Video is 4x real time.

